# Super homotypic targeting by exosome surface engineering

**DOI:** 10.1101/2024.05.01.592036

**Authors:** Huai-Song Wang, Tianben Ding, Yuhong Liu, Yuqi Zhou, Yaqi Zhao, Mika Hayashi, Xin-Yuan Hu, Zi-Wei Yang, Natsumi Tiffany Ishii, Hiroki Matsumura, Anel Umirbaeva, Hongwei Guo, Jing-Lian Su, Yin-Yu Yan, Fu-Han Gao, Jia-Jing Li, Nao Nitta, Masako Nishikawa, Yutaka Yatomi, Ya Ding, Masahiro Sonoshita, Dino Di Carlo, Shiro Suetsugu, Keisuke Goda

## Abstract

Homotypic targeting is the inherent ability of cells for preferential interaction with cells of similar or identical types, a phenomenon commonly seen in cell adhesion, tissue formation, and immune responses. Unfortunately, its full potential remains largely untapped. Here we introduce an approach to drastically boost the homotypic targeting capabilities of cells via exosomes (nanoscale extracellular vesicles secreted by cells). By engineering exosome surfaces with lanthanides, we amplify specific cell-exosome interactions by more than 25-fold, significantly accelerating the selective capture of exosomes by cells of the same lineage. This substantial enhancement in cellular homophilicity opens up an entirely new class of applications, two of which we showcase here with unprecedented performance: using cells to detect specific exosomes and using exosomes to detect specific cells. The concept of “super homotypic targeting” offers enormous potential to transform cancer diagnostics, immunotherapy, targeted drug delivery, tissue engineering, and vaccine development.

---

Homotypic targeting is a fundamental biological concept where cells preferentially interact or bind with cells of similar or identical types^1–5^. This homophilic specificity is crucial in various physiological processes, including cell adhesion, tissue formation, and immune responses^6–9^. In therapeutic development, especially in targeted drug delivery, homotypic targeting is important in ensuring that drugs are delivered primarily to target cells, reducing systemic side effects and increasing treatment efficacy^5,10–13^. Furthermore, in tissue engineering, homotypic targeting facilitates the controlled interaction and formation of tissues by cells^14,15^. In the field of immunology, it is pivotal in enabling T cells to engage in homotypic interactions during the formation of immunological synapses, which are essential for mounting effective immune responses^16,17^. Nonetheless, the full potential of homotypic targeting remains largely untapped, mainly due to challenges in achieving high specificity, fast responses, and preventing off-target effects in complex and heterogenous conditions such as cancer^1,10,12,13^.

In this study, we introduce an approach to drastically boost the homotypic targeting capabilities of cells via exosomes (small extracellular vesicles secreted by cells, typically ranging from 30 to 150 nm in diameter), achieved by engineering the surface of exosomes with lanthanide ions such as Eu^3+^ and Tb^3+^ (Fig. 1a). We leverage the natural overexpression of sialic acid (SA) on cell membranes, a common characteristic in cancer cells^18–20^, for competitive coordination with these lanthanide ions. This unique coordination amplifies specific cell-exosome interactions by more than 25-fold, significantly speeding up the selective capture of exosomes by cells of the same lineage, even in the excessive presence of other exosomal populations (Fig. 1a). This amplification, which we refer to as “super homotypic targeting”, is not just a marginal improvement but a substantial enhancement in homotypic targeting efficiency. It reduces the exosome capture time of the cells from over 2 days to only about 3 hours, offering a distinct strategy for discriminating exosomes from different origins in a highly heterogeneous population of exosomes (Fig. 1b). In other words, cells can act as selective instruments for distinguishing between various exosome types, based on their capture speed. Consequently, this substantial enhancement in cellular homophilicity opens up an entirely new class of applications, two notable examples of which we showcase here with unprecedented performance: (i) using cells to detect specific exosomes; (ii) using exosomes to detect specific cells. The concept of “super homotypic targeting” holds immense potential for revolutionizing cancer diagnostics, immunotherapy, targeted drug delivery, tissue engineering, and vaccine development.

**Fig. 1.**
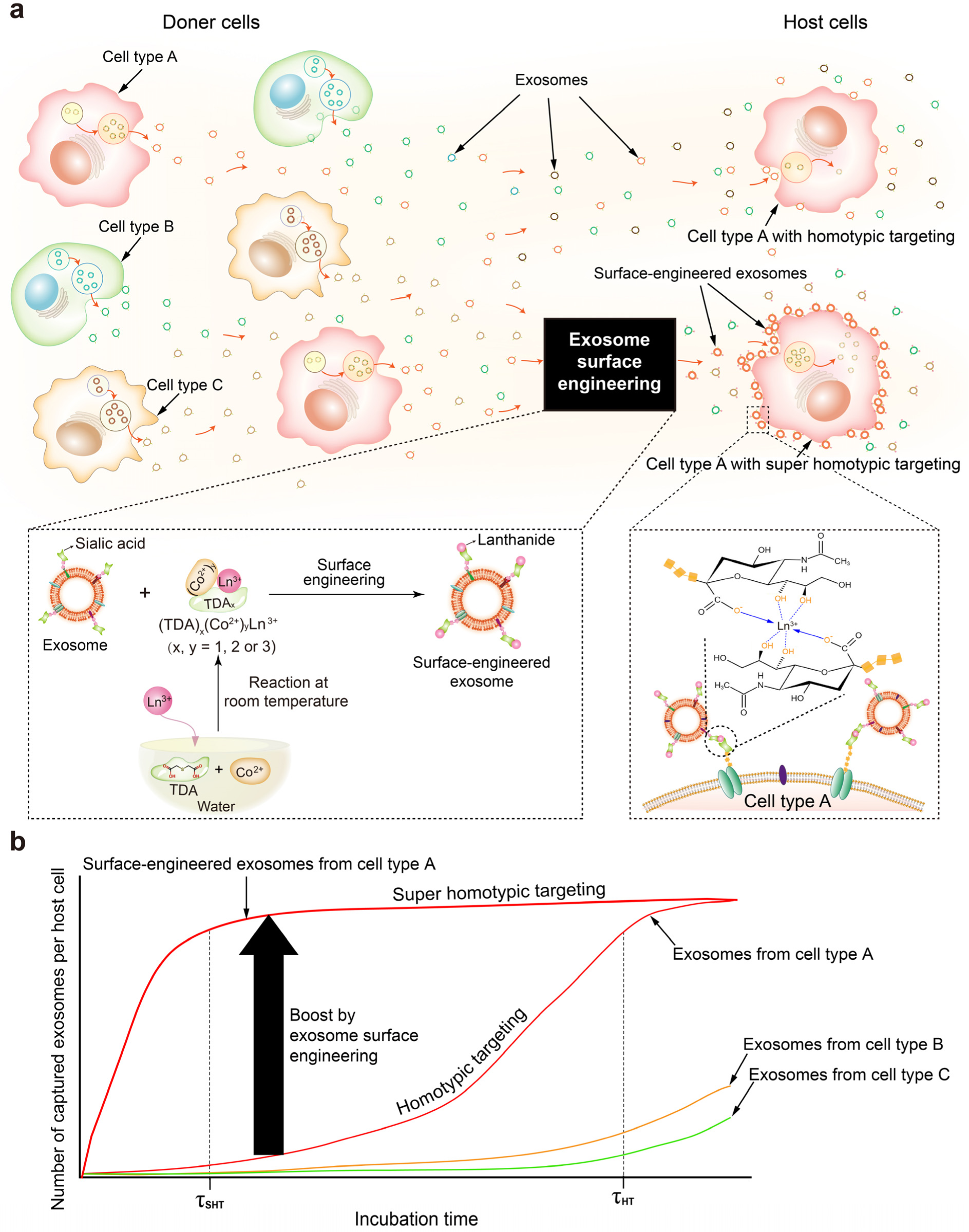
Concept of super homotypic targeting. **a**, Schematic. The homotypic targeting capabilities of host cells (type A) are significantly boosted by engineering exosome surfaces with lanthanide ions. **b,** Exosome capture rates of host cells (type A) when incubated in a pool of exosomes from various donor cells (types A-C). Exosome surface engineering amplifies specific cell-exosome interactions, thereby significantly speeding up the selective capture of exosomes by cells of the same lineage (type A), even in the excessive presence of exosomes from other cell types (type B, type C). This amplification dramatically reduces the exosome capture time of the host cells ( *τ*_SHT_ ≪ *τ*_HT_), offering a distinct strategy for discriminating exosomes from different origins in the heterogeneous population of exosomes. In other words, host cells can act as selective instruments for distinguishing between various exosome types, based on their capture speed.

## Exosome surface engineering

To experimentally demonstrate super homotypic targeting by surface engineering of exosomes, we focused on the selective capture of exosomes derived from MDA-MB-436 triple-negative breast cancer (TNBC) cells (corresponding to cell type A in Fig. 1a) in the excessive presence of exosomes derived from MCF-7 breast cancer (BC) cells and blood cells (corresponding to cell types B and C, respectively, in Fig. 1a). TNBC is the most challenging subtype of BC to detect due to its absence of the three most common types of BC cell surface receptors (ER, PR, HER2) despite its notorious recurrence and aggressive nature^21^. We targeted SA, primarily *N*-acetylneuraminic acid, a key molecule in cancer malignancy and metastasis^18–20^. The bimetallic compound TDA-Co-Eu, composed of 2,2’-thiodiacetic acid (TDA), Eu^3+^, and Co^2+^, was utilized for its selective affinity to SA on cancer cell surfaces^22^. This unique capability arises from the competitive binding between SA and Eu^3+^ that is released from the complex. Our preparation of TDA-Co-Eu was streamlined by sequentially mixing solutions of TDA, Co^2+^, and Eu^3+^ (Fig. 1a inset). Mass spectrometry revealed a mixture of compounds, including (TDA)Co^2+^Eu^3+^, (TDA)(Co^2+^)_2_Eu^3+^, (TDA)_2_(Co^2+^)_2_Eu^3+^, and (TDA)_3_(Co^2+^)_2_Eu^3+^ (Fig. 2a). Despite being a compound mix, the (TDA)_x_(Co^2+^)_y_Eu^3+^ complex remains effective for SA sensing, as demonstrated by the detected (SA)_2_Eu^3+^ in the mass spectrum post SA reaction (Fig. 2b). Fluorometric analysis further validated the ability of SA to liberate Eu^3+^ ions from (TDA)_x_(Co^2+^)_y_Eu^3+^, as indicated by the observable Eu^3+^ fluorescence that is typically inhibited by Co^2+^ (Fig. 2c). Notably, the (TDA)_x_(Co^2+^)_y_Eu^3+^ complex displayed a pronounced (over 10-fold) sensitivity to SA over other biomolecules (Fig. 2d). Subsequently, exosome surfaces were modified using Eu^3+^ ions, introducing the (TDA)_x_(Co^2+^)_y_Eu^3+^ solution to exosome suspensions followed by ultracentrifugation to remove unreacted Eu^3+^ ions and residual (TDA)_x_(Co^2+^)_y_. Exosomes from MDA-MB-436 cells, MCF-7 cells, and blood cells present in FBS, termed Exo-MDA, Exo-MCF7, and Exo-BLD (Extended Data Fig. 1a-d), underwent this Eu^3+^ ion-based surface modification. Inductively coupled plasma mass spectrometry (ICP-MS) highlighted the Eu^3+^ content in these modified exosomes, coined Exo-MDA-Eu, Exo-MCF7-Eu, and Exo-BLD-Eu (Fig. 2e). The increased content in Exo-MDA-Eu exosomes is likely attributed to the prominent SA expression on MDA-MB-436 cell membranes (Fig. 2f), causing Exo-MDA to possess a higher SA concentration, which then binds more Eu^3+^ from (TDA)_x_(Co^2+^)_y_Eu^3+^. The rise in zeta potentials of these surface-engineered exosomes, compared to their unaltered counterparts, highlights the effective incorporation of Eu^3+^ (Fig. 2g).

**Fig. 2.**
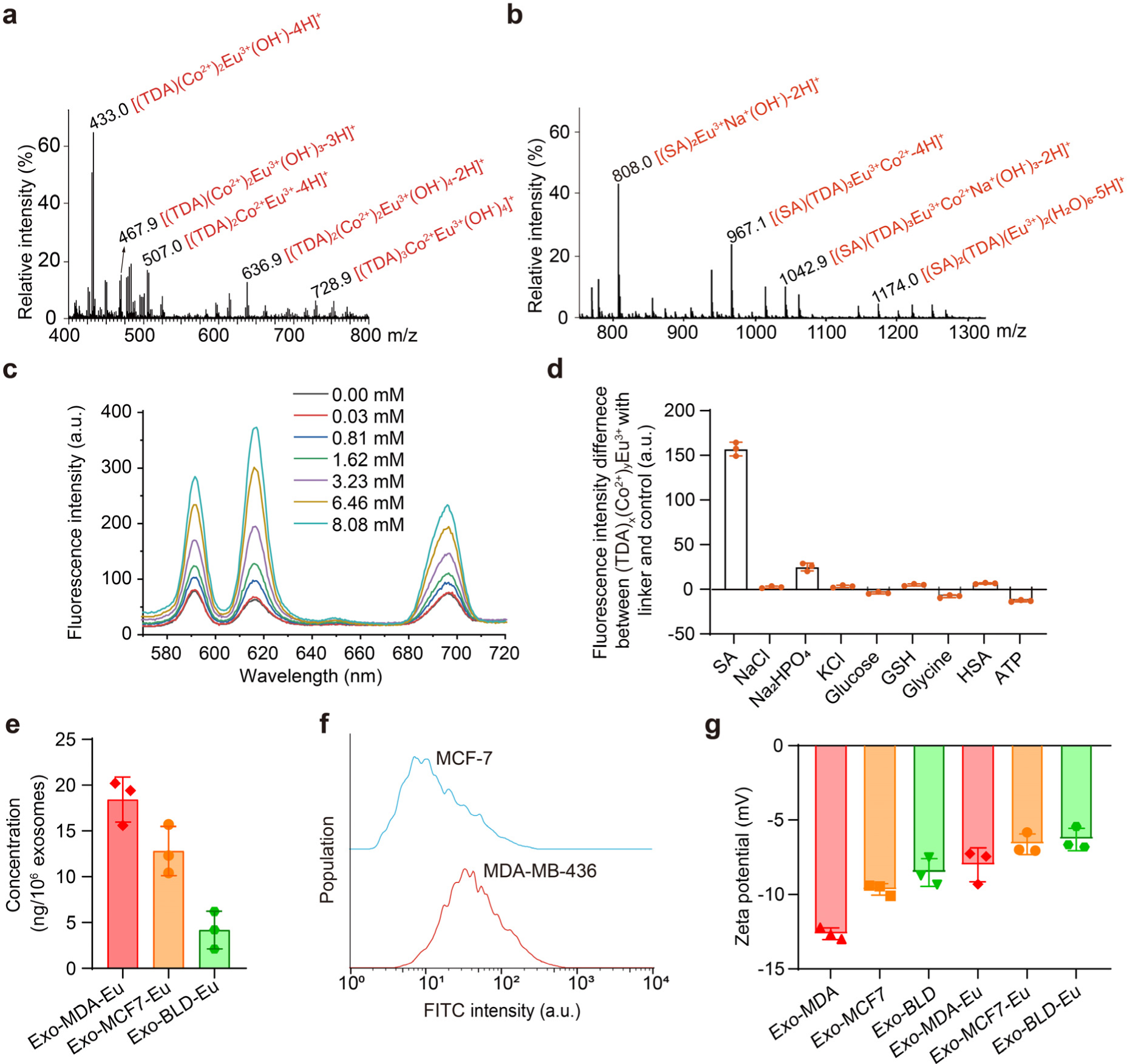
Exosome surface modification with Eu^3+^ ions for super homotypic targeting. **a**, Mass spectrum of (TDA)_x_(Co^2+^)_y_Eu^3+^. **b,** Mass spectrum of (TDA)_x_(Co^2+^)_y_Eu^3+^ after its reaction with SA. **c,** Fluorescence spectra of (TDA)_x_(Co^2+^)_y_Eu^3+^ for sensing different concentrations of SA (excitation wavelength: 390 nm). **d,** Fluorescence intensity differences between (TDA)_x_(Co^2+^)_y_Eu^3+^ with different linkers and the control. Error bars represent standard deviations (n = 3). **e,** ICP-MS analysis of Eu^3+^ content in different surface-engineered exosomes. **f,** Flow-cytometric enumeration of MCF-7 and MDA-MB-436 cells labeled with FITC-SNA. **g,** Zeta potentials of surface-engineered and -unengineered (intact) exosomes. Error bars represent standard deviations (n = 3).

Elevated SA levels facilitated Eu^3+^ ion binding to host cancer cell and exosome membranes. Each Eu^3+^ ion coordinated with two SA molecules and thus acted as enhancers of exosome-mediated homotypic targeting (Fig. 1a inset). Laser scanning confocal microscopy (LSCM) of Eu^3+^-modified exosomes (Exo-MDA-Eu, Exo-MCF7-Eu, Exo-BLD-Eu), labeled with PKH67 (green), highlighted that Exo-MCF7-Eu exosomes were incorporated by MDA-MB-436 cells after 12 h, with the fluorescence intensity approaching an appreciable level at 20 h (Fig. 3a, b). In contrast, Exo-MDA-Eu achieved similar fluorescence intensity in only 6 h with MDA-MB-436 cells, indicating its quicker targeting capability (Fig. 3c). The results validated the improved exosome-cell interaction after the surface modification in comparison to exosomes without the surface modification (Fig. 3c and Extended Data Fig. 1e, f). Enhanced interactions with MCF-7 cells were observed for both Exo-MDA-Eu and Exo-MCF7-Eu exosomes compared to their non-modified counterparts (Extended Data Fig. 1f, g). These findings highlight the role of Eu^3+^ ions in boosting the exosome-mediated homotypic targeting efficiency of cells. In a subsequent analysis, we measured the PKH67 fluorescence, which signals exosome capture, in MDA-MB-436 and MCF-7 cells over different incubation durations. Notably, MCF-7 cells exhibited a pronounced capture of Exo-MDA-Eu exosomes compared to Exo-MCF7-Eu (Extended Data Fig. 1h, i), which is presumably attributed to a denser Eu^3+^ ion presence in the former. Further, PKH67 fluorescence tracking revealed that host MDA-MB-436 cells preferentially absorbed Exo-MDA-Eu exosomes, with substantially faster capture than other exosome types (Fig. 3a-i and Extended Data Fig. 1f, j). At the 3rd hour, the homotypic targeting efficiency of host MDA-MB-436 cells had increased by a remarkable 25.6-fold due to the application of Exo-MDA-Eu exosomes Fig. 3d). LSCM confirmed that Exo-MDA-Eu uniformly populated MDA-MB-436 cells after a 3-h incubation, whereas Exo-MCF7-Eu and Exo-BLD-Eu exosomes required 12 h and 40 h, respectively (Fig. 3b, c, e-h). Increasing exosome concentration reduced capture time, demonstrating the host cells’ capacity to differentiate between exosome types based on absorption rates (Fig. 3j-l). Thus, MDA-MB-436 cells emerged as selective tools for discerning between the three exosome types based on rapid, medium, and slow capture timelines, outpacing unmodified exosomes. This temporal differentiation offered a strategy for exosome discrimination and the 3-h specific incubation period was chosen to maximize specificity in detecting Exo-MDA exosomes from a mixture of Exo-MDA, Exo-MCF7, and Exo-BLD exosomes.

**Fig. 3.**
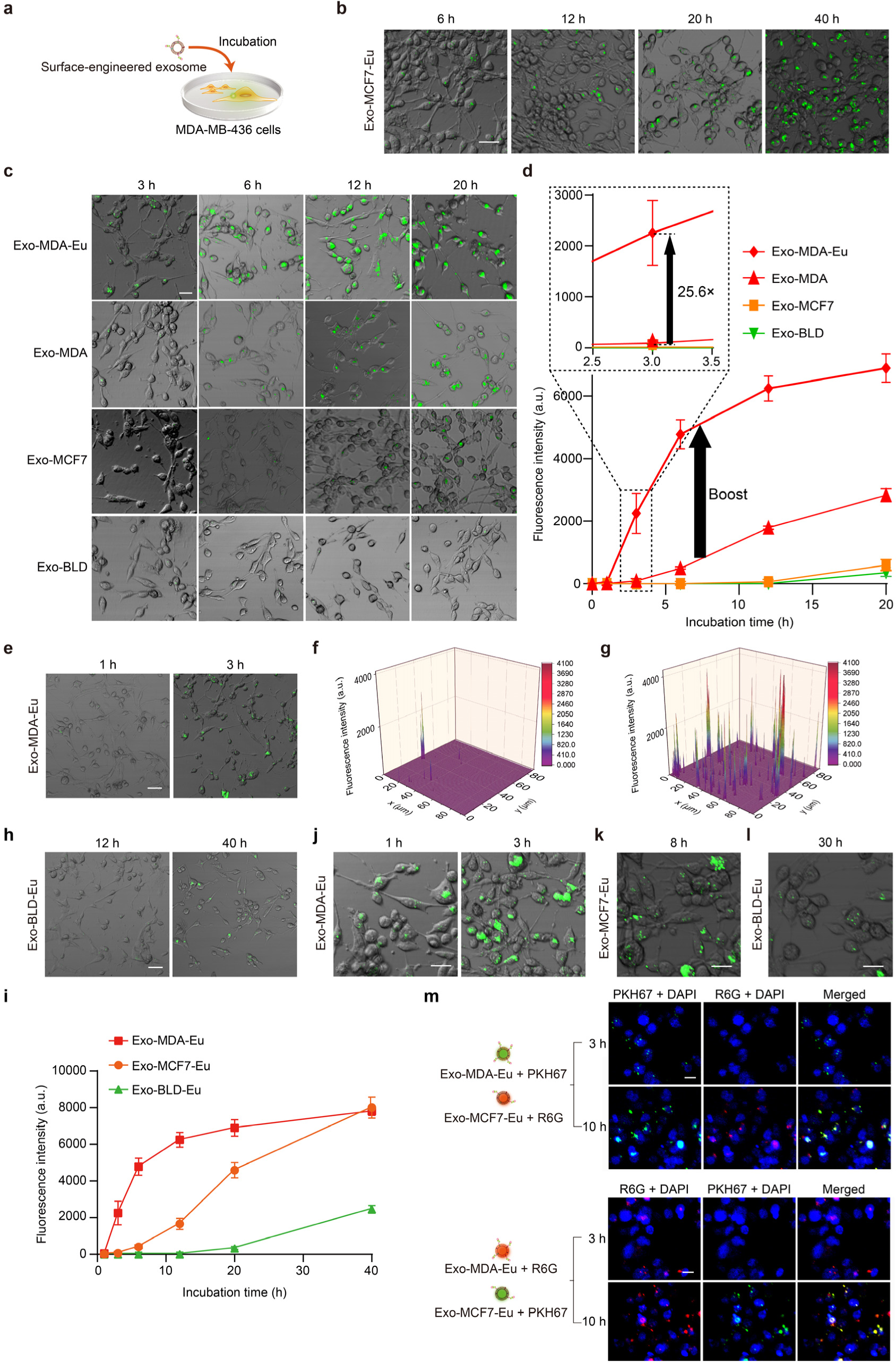
LSCM images of cancer cells incubated with surface-engineered exosomes. **a**, Schematic of mixing MDA-MB-436 cells with various surface-engineered exosomes. **b,** LSCM-based monitoring of MDA-MB-436 cells incubated with Exo-MCF7-Eu. Exosomes were labeled with PKH67 (green). Scale bars: 30 μm. **c,** Merged bright-field and fluorescence images of host MDA-MB-436 cells after incubation with exosomes from donor MDA-MB-436 cells (Exo-MDA-Eu) via exosome surface engineering, exosomes from donor MDA-MB-436 cells (Exo-MDA), exosomes from donor MCF-7 cells (Exo-MCF7), and exosomes from donor blood cells (Exo-BLD) for 3, 6, 12, and 20 h. Scale bar: 30 μm. **d,** Intensity of fluorescence emitted from the captured exosomes per host MDA-MB-436 cell varied over the course of the incubation period. After a 3-h incubation, the host MDA-MB-436 cells exhibited a significantly enhanced efficiency in capturing Exo-MDA-Eu exosomes; 25.6, 834, and 2503 times higher than with Exo-MDA, Exo-MCF7, and Exo-BLD exosomes, respectively. **e,** LSCM-based monitoring of MDA-MB-436 cells incubated with Exo-MDA-Eu for 1 h and 3 h. Scale bar: 50 μm. **f, g,** Fluorescence intensity values of Exo-MDA-Eu in MDA-MB-436 cells after 1-h (**f**) and 3-h (**g**) incubation. **h,** LSCM-based monitoring of MDA-MB-436 cells incubated with Exo-BLD-Eu for 12 and 40 h. The concentration of the Exo-BLD-Eu suspension is ∼3.8×10^8^ exosomes/mL. Scale bars: 50 μm. **i,** Surface-engineered exosome capture for Exo-MCF7-Eu, Exo-MDA-Eu, and Exo-BLD-Eu by MDA-MB-436 cells. Error bars represent standard deviations (n = 3). **j-l,** MDA-MB-436 cells incubated with 40 μL suspensions of Exo-MDA-Eu (**j**, ∼4.0×10^7^ exosomes/mL), Exo-MCF7-Eu (**k**, ∼4.0×10^7^ exosomes/mL), and Exo-BLD-Eu (**l**, ∼3.8×10^8^ exosomes/mL). By increasing the concentration of exosomes, MDA-MB-436 cells were able to capture Exo-MDA-Eu within 1 h, Exo-MCF7-Eu within 8 h, and Exo-BLD-Eu within 30 h. Scale bars: 40 μm. **m,** LSCM images of host MDA-MB-436 cells incubated with Exo-MDA-Eu and Exo-MCF7-Eu exosomes for 3 h and 10 h, respectively. Top: Exo-MDA-Eu labeled with PKH67 and Exo-MCF7-Eu labeled with R6G. Bottom: Exo-MDA-Eu labeled with R6G and Exo-MCF7-Eu labeled with PKH67. The cell nuclei were stained with DAPI (blue). Scale bars: 30 μm.

To rigorously validate the capacity of MDA-MB-436 cells for capturing exosomes derived from other MDA-MB-436 cells that simulate exosomes derived from TNBC cells, we employed two-color intracellular fluorescence imaging. In our two-color fluorescence imaging, Exo-MDA-Eu and Exo-MCF7-Eu exosomes were distinctly tagged with PKH67 (green) and Rhodamine 6G (R6G, red). Their interaction with MDA-MB-436 cells was monitored. By 3-h incubation, fluorescence predominantly traced Exo-MDA-Eu exosome capture, regardless of the fluorophore used (Fig. 3m). However, by 10-h incubation, dual fluorescence denoted the presence of both exosome types in MDA-MB-436 cells, underscoring the preferential affinity of MDA-MB-436 cells for Exo-MDA-Eu exosomes. *In vivo* imaging further elucidated the robust interaction of Exo-MDA-Eu exosomes with TNBC cells^23,24^. Both Exo-MDA-Eu and Exo-MDA exosomes were tagged with the near-infrared fluorescence probe, DiR. Following inoculation of MDA-MB-436 cells into BALB/c nude mice and tumor emergence at the axillary region (7 days after inoculation), either Exo-MDA-Eu or Exo-MDA exosomes (DiR-tagged) were administered via intratumoral or tail vein injections (Extended Data Fig. 2a-c). Imaging at varied intervals revealed prolonged tumor visibility for Exo-MDA-Eu exosomes versus non-Eu^3+^-labeled and pure DiR controls (Extended Data Fig. 2d). Notably, tumor fluorescence peaked within 24 h (Extended Data Fig. 2e). *Ex vivo* imaging affirmed this, with tumors radiating pronounced fluorescence relative to other organs (Extended Data Figs. 2f-h). Limited detection of Exo-MDA-Eu exosomes in the liver and kidneys suggests preferential tumor localization and low biodistribution. These results highlight Exo-MDA-Eu’s potential in tumor visualization and drug delivery support. Interestingly, after intravenous administration, we noted substantial Exo-MDA-Eu exosome deposition in the spleen (Extended Data Fig. 2i-k), consistent with prior reports^25–27^. This could be attributed to the immunological role of the spleen, with resident macrophages poised for exosome clearance, leading to Exo-MDA-Eu exosome accumulation.

## Extreme-precision exosome detection based on super homotypic targeting

Exosomes carry proteins, nucleic acids, and lipids, reflecting the state of their origin cells, and are stable in body fluids, making them potential biomarkers for diseases such as cancer^28–31^. Exosomes are key in liquid biopsies for non-invasive cancer detection, providing insights into cancer progression and helping monitor treatment responses^32–34^. While numerous methods have been developed to detect exosomes in a blood or serum sample, their specificity, however, falls short in differentiating tumor-specific exosomes from the overall exosomal population in the sample^35,36^. Consequently, these methods tend to focus on analyzing specific proteins or nucleic acids within the entire sample. The shared characteristics of proteins and nucleic acids within different exosomes complicate their analysis^37,38^. We address these problems with super homotypic targeting.

Our extreme-precision exosome detector based on super homotypic targeting is schematically shown in Fig. 4a. Here, we leveraged MDA-MB-436 cells as host cells to selectively capture and enrich exosomes derived from TNBC tumor cells (Exo-TNBC) in the excessive presence of other exosomal populations from BC cells and blood cells (Exo-MCF7, Exo-BLD, respectively) and subsequently analyzed the enriched exosomes on the cells using flow cytometry. Specifically, a blood sample was collected from an animal subject (a mouse in this study) and subjected to centrifugation and ultracentrifugation to extract a pool of Exo-TNBC, Exo-BC, and Exo-BLD exosomes. Exo-TNBC exosomes were modified with Eu^3+^ ions and fluorescently labeled with PKH67 (Extended Data Fig. 3a). During a 3-h incubation, only Exo-TNBC exosomes were captured and enriched by MDA-MB-436 cells (host cells), which were measured for accurate enumeration with a commercially affordable flow cytometer (BD FACSCelesta) and a home-made imaging flow cytometer^39,40^. As mentioned above, this incubation time (3 h) was carefully chosen to optimize the specificity of detecting Exo-TNBC exosomes from the heterogeneous exosome population (Fig 3i, 4b-e and Extended Data Fig. 3b-g).

**Fig. 4.**
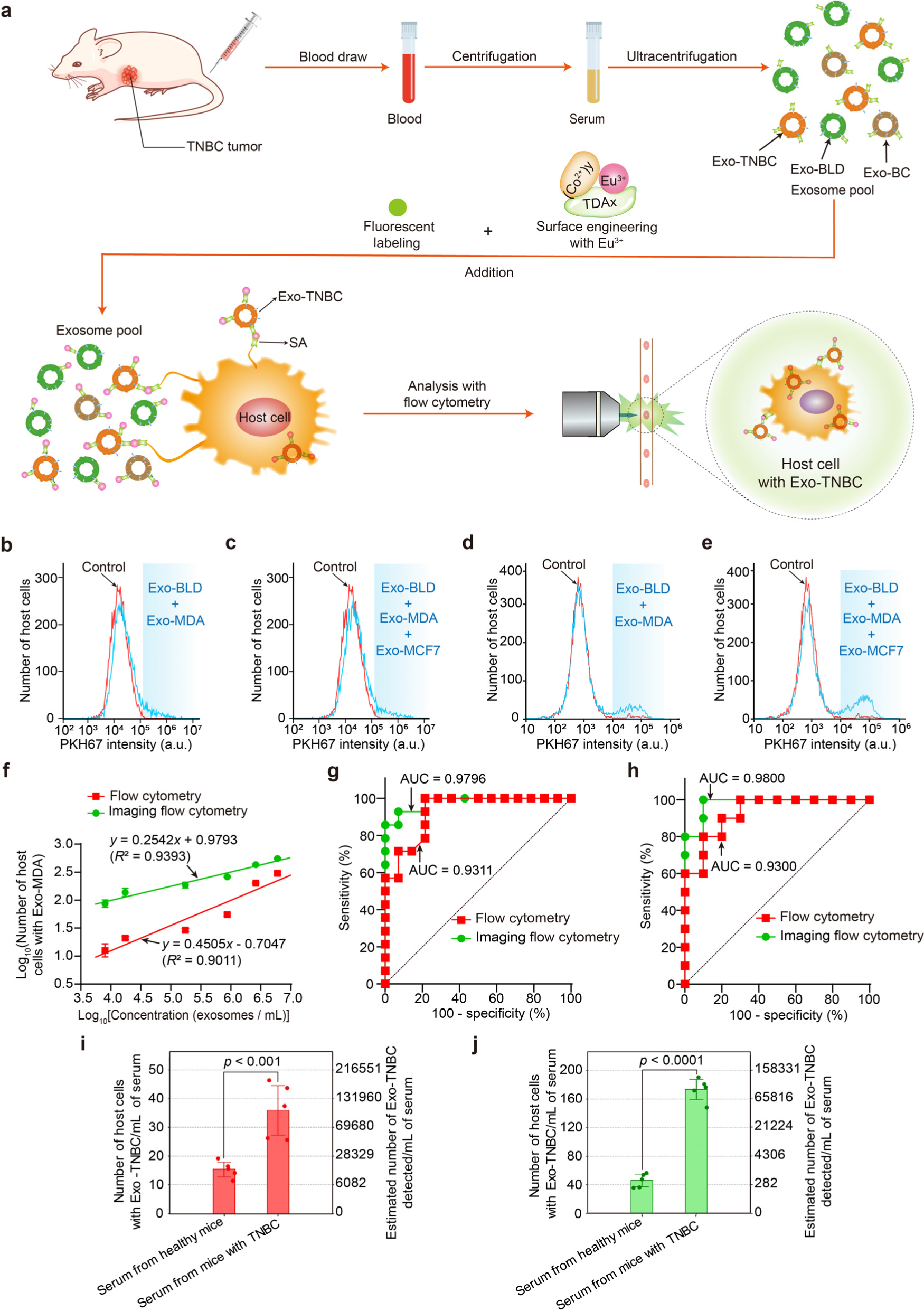
Schematic and results of extreme-precision exosome detection based on super homotypic targeting. **a**, Schematic. A blood sample is collected from a mouse and subjected to centrifugation and ultracentrifugation to extract Exo-TNBC, Exo-BC, and Exo-BLD exosomes. Exosomes are modified with Eu^3+^ ions and fluorescently labeled with PKH67. During a 3-h incubation, only Exo-TNBC exosomes are captured and enriched by MDA-MB-436 cells (host cells), which are measured for accurate enumeration with a commercially affordable flow cytometer and a home-made imaging flow cytometer. **b, c,** Flow cytometry of host cells selectively capturing Exo-MDA in serum (1 mL) containing 2×10^5^ Exo-MDA exosomes and 4×10^8^ Exo-BLD exosomes without (**b**) and with (**c**) 5×10^7^ Exo-MCF7 exosomes, in comparison with the control (host cells without incubation with the exosomes). 10,000 host cells were used for each measurement. Light-blue regions indicate counting thresholds. **d, e,** Imaging flow cytometry of host cells selectively capturing Exo-MDA in serum (1 mL) containing 2×10^5^ Exo-MDA exosomes and 4×10^8^ Exo-BLD exosomes without (**d**) and with (**e**) 5×10^7^ Exo-MCF7 exosomes, in comparison with the control (host cells without incubation with the exosomes). 10,000 host cells were used for each measurement. Light-blue regions indicate counting thresholds. **f,** Number of host cells selectively capturing Exo-MDA exosomes derived from MDA-MB-436 cells at different spiked concentrations in serum (1 mL) containing about 4×10^8^ Exo-BLD exosomes derived from blood cells, detected with flow cytometry and imaging flow cytometry. Error bars represent standard deviations (n = 3). **g,** ROC curves for detecting Exo-MDA exosomes in the presence of Exo-BLD exomes. **h,** ROC curves for detecting Exo-MDA exosomes in the presence of Exo-BLD and Exo-MCF7 exosomes. **i,** Number of host cells selectively capturing TNBC-derived exosomes (Exo-TNBC) in the serum of TNBC-afflicted mice, measured with flow cytometry. The result is compared with the control (healthy mice): n = 5 for each category. 3,000 host cells were used for each measurement. An axis for the estimated number of detected exosomes based on the linear fit in **f** is also shown on the right. The *p*-value was obtained using a two-sided t-test to assess the statistical significance of the difference between the two groups. **j,** Number of host cells selectively capturing Exo-TNBC exosomes in the serum of TNBC-afflicted mice, measured with imaging flow cytometry. The result is compared with the control (healthy mice): n = 5 for each category. 3,000 host cells were used for each measurement. An axis for the estimated number of detected exosomes based on the linear fit in **f** is also shown on the right. The *p*-value was obtained using a two-sided *t*-test to assess the statistical significance of the difference between the two groups.

To evaluate the sensitivity, specificity, and limit of detection (LOD) of the exosome detector, we performed spike-and-recovery tests with Exo-MDA exosomes (1×10^4^–1×10^7^ exosomes/mL) and/or Exo-MCF7 exosomes (5×10^7^ exosomes/mL) spiked into 1.0 mL of fetal bovine serum (FBS) (4×10^8^ exosomes/mL) to mimic serum samples laden with Exo-TNBC exosomes at pathologically relevant concentrations. Flow cytometry results revealed a signal intensity boost ranging from 0.75% to 23.03%, correlating with Exo-MDA exosome concentrations between 1.74×10^4^ and 1.04×10^7^ exosomes/mL (Extended Data Fig. 3d, h). These results highlight the affinity of the host cells for Exo-MDA exosomes, which persisted even in the presence of Exo-MCF7 exosomes (2.0×10^5^ exosomes/mL) (Extended Data Fig. 3e-g). A compelling observation was the strong linear relationship when plotting the Exo-MDA exosome concentration logarithmically against the count of the host cells containing Exo-MDA exosomes, indicated by a linear fit with R^2^ = 0.9011 (Fig. 4f). Based on the linear fit, the LOD was estimated to be approximately 751 exosomes/mL of serum containing 4×10^8^ Exo-BLD exosomes, using the formula *LOD* = 3.3*σ*/*S*, where *σ* is the standard deviation of the y-intercept and *S* is the slope of the linear fit^41^. Furthermore, Fig. 4g shows exceptional sensitivity and specificity for the detection of Exo-MDA exosomes in the serum sample (1 mL), even including Exo-MCF7 exosomes (Fig. 4h), as evidenced by the receiver operating characteristic (ROC) curve boasting an area-under-the-curve (AUC) value of 0.9300 for the selective detection of Exo-MDA exosomes in the presence of Exo-BLD and Exo-MCF7 exosomes (see similar results in Extended Data Fig. 4a, b that show the specific detection of Exo-MDA in the presence of Exo-BLD with leukemia-derived exosomes). The imaging flow cytometry showed a marked improvement in both sensitivity and specificity compared with the traditional flow cytometry (Fig. 4d, e and Extended Data Fig. 4c) because it effectively eliminated noisy background fluorescence signals to accurately pinpoint false positive events that arose from cell death or debris^39,42^ (Extended Data Fig. 4d-f). As a result of these enhancements, our data analysis revealed a stronger linear correlation with Exo-MDA exosome concentrations, highlighted by a notably high coefficient of determination (R^2^ = 0.9393, Fig. 4f). Additionally, we observed a 10-fold boost in the estimated LOD value, settling at 63 exosomes/mL of serum containing 4×10^8^ Exo-BLD exosomes. The ROC curve was also pronounced, with an AUC value of 0.9796 (Fig. 4g), even including Exo-MCF7 exosomes (Fig. 4h).

We employed the fully characterized exosome detector to directly identify Exo-TNBC exosomes in serum samples from TNBC-afflicted mice (Methods). After 14 days of TNBC growth in the mice, blood samples were collected. 1 mL of serum was isolated from the blood samples by centrifugation. The serum samples were analyzed with both the flow cytometer and imaging flow cytometer. Results show that flow cytometry effectively differentiated between serum samples of healthy mice (n = 5) and the TNBC tumor-bearing mice (n = 5) with a high statistical significance (*p* < 0.001) by identifying TNBC-derived exosomes (Fig. 4i). When using imaging flow cytometry, as depicted in Fig. 4j, the distinction between the groups became even clearer, reflected by an even lower *p*-value (*p* < 0.0001). These excellent *p*-values are consistent with the aforementioned LODs and ROC curves in Fig. 4g, h.

## Extreme-precision CTC detection based on super homotypic targeting

Circulating tumor cells (CTCs), which are cancer cells shed from primary tumors into the bloodstream, serve as vital indicators of the potential spread of cancer and can be early markers of metastasis, thereby aiding in the development of liquid biopsies^43–45^. Their detection is crucial for cancer diagnosis but presents significant challenges due to their sparse presence among the vast, diverse populations of blood cells^46,47^. Several factors compound these difficulties: the absence of specific biomarkers for reliable CTC identification, the need for large sample volumes to capture even a single CTC or to gather statistically significant CTC populations, and the phenotypic variability that CTCs exhibit, depending on their origin^48–51^. Many traditional detection methods lack the required sensitivity, specificity, or both. While conventional flow cytometry can detect CTCs, it requires time-consuming pre-enrichment steps and multiple sorting runs^48,52^. Specialized techniques that can identify CTCs are often costly and require intricate setups^48,53^, limiting their widespread application. We address these limitations with super homotypic targeting.

Our extreme-precision CTC detector based on super homotypic targeting is schematically shown in Fig. 5a. Here, exosomes were leveraged as signal amplifiers for the ultrasensitive detection of rare CTCs with single-cell resolution in a single run of standard flow cytometry. By enhancing the homotypic targeting capabilities of CTCs via engineering the surface of Exo-TNBC exosomes, this method employed probes specifically directed to MDA-MB-436 cells and produced fluorescence from approximately 10,000 MDA-MB-436 cells (reporter cells) for each CTC, effectively boosting the flow cytometer’s ability to detect them. Essentially, this method magnified the apparent number of rare CTCs by about four orders of magnitude, echoing the function of operational amplifiers in electronics. Specifically, a blood sample was collected from an animal subject (a mouse in this study) with TNBC and subjected to centrifugation for isolating TNBC-derived CTCs, CTCs derived from other cancer types, and peripheral blood mononuclear cells (PBMCs). The surface of Exo-TNBC was modified with Tb^3+^ ions released from the metal complex TDA-Co-Tb (TCT) comprising TDA, Co^2+^, and Tb^3+^ ions^22^, employing a procedure similar to the established protocol used for the Eu^3+^ ion surface modification (Methods), after loading the exosomes (Exo-MDA) with hexaminolevulinate (HAL)^50,54,55^. The HAL-loaded surface-engineered exosomes were added to the blood sample and incubated with it such that they could specifically bind to the CTCs. After removing uncaptured exosomes, all CTCs and PBMCs were completely disrupted by ultrasonication, resulting in the release of HAL. The released HAL was incubated with 2×10^6^ reporter cells to convert HAL into the fluorescent compound, protoporphyrin IX (PpIX), within the cells, where PpIX produced strong fluorescence in mitochondria due to their reduced metabolism in converting PpIX to heme^55^ (Fig. 5b). The reporter cells were analyzed with an affordable flow cytometer operating in single-color mode (BD FACSCelesta) to count the fluorescent population of the reporter cells in a single measurement run.

**Fig. 5.**
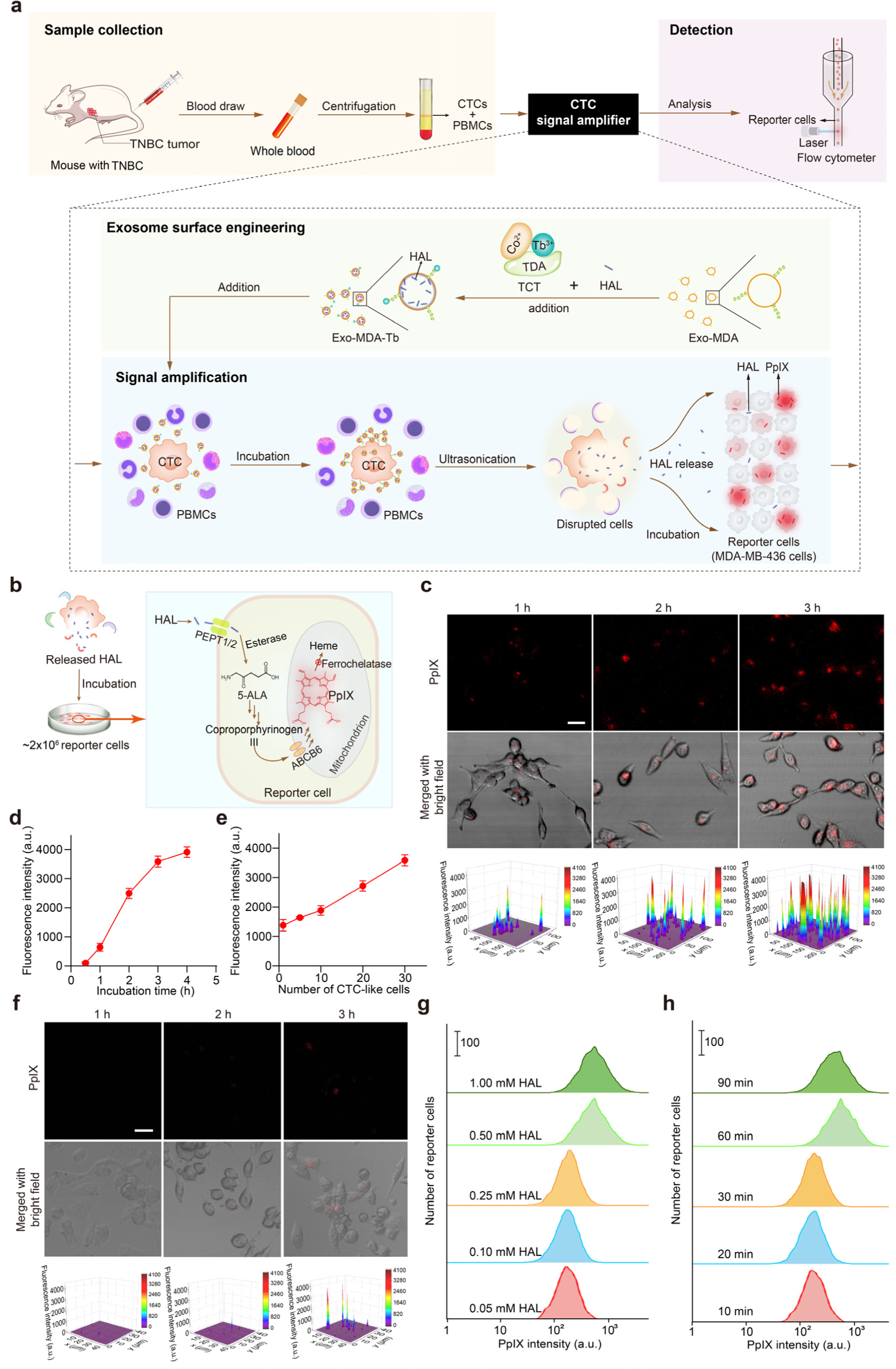
Schematic of extreme-precision CTC detection and validation of signal amplification of CTC-like cells based on super homotypic targeting. **a**, Schematic. The detection procedure is as follows: (i) a blood sample is collected from a mouse with TNBC and subjected to centrifugation for isolating TNBC-derived CTCs, other CTCs, and PBMCs; (ii) the surface of exosomes derived from cells of the MDA-MB-436 TNBC cell line is engineered with Tb^3+^ ions, after loading the exosomes with HAL; (iii) the HAL-loaded surface-engineered exosomes are added to the blood sample and incubated with it so that they can specifically bind to the TNBC-derived CTCs; (iv) after removing uncaptured exosomes, all CTCs and PBMCs are completely disrupted by ultrasonication, resulting the release of HAL; (v) the released HAL is incubated with reporter cells (MDA-MB-436 cells) to convert HAL into PpIX within the cells; (vi) the reporter cells are analyzed with a flow cytometer to count the fluorescent population of the reporter cells. **b,** Schematic of staining MDA-MB-436 cells (reporter cells) with HAL released by ultrasonicated CTC-like cells and turning them into reporter cells. **c,** LSCM images and fluorescence intensities of reporter cells after incubation with ultrasonicated PBMCs with 30 spiked CTC-like cells for varying incubation durations (1-3 h). The PBMCs with CTC-like cells were pre-incubated with HAL-loaded Exo-MDA-Tb (1.6×10^6^ exosomes/mL) in a culture medium with 0.8 µg/mL TCT for 1 h. Scale bar: 20 μm. **d,** Fluorescence intensity of suspended reporter cells after incubation with ultrasonicated PBMCs with 30 spiked CTC-like cells for varying incubation durations (0.5-4 h). Error bars represent standard deviations (n = 3). Fluorescence intensity values shown in the figure were obtained by calculating the total fluorescence intensity from each cell and dividing it by the total number of cells (n = 30-50). **e,** Fluorescence intensity of suspended reporter cells after a 3-h incubation with ultrasonicated PBMCs with varying spiked CTC-like cell numbers (1-30 cells). Error bars represent standard deviations (n = 3). Fluorescence intensity values shown in the figure were obtained by calculating the total fluorescence intensity from each cell and dividing it by the total number of cells (n = 30-50). **f,** LSCM images and fluorescence intensities of reporter cells after incubation with ultrasonicated PBMCs with a single spiked CTC-like cell for varying incubation durations (1-3 h). Scale bars: 20 μm. **g,** Influence of the HAL concentration on flow cytometry of reporter cells. Exo-MDA-Tb was incubated with varying HAL concentrations (0.05-1.00 mM) for 1 h before incubation with 30 CTC-like cells and HAL introduction to reporter cells. **h,** Influence of the HAL incubation duration on flow cytometry of reporter cells. Exo-MDA-Tb was incubated with 0.5 mM HAL for varying incubation durations (10-90 min) before incubation with 30 CTC-like cells and HAL introduction to reporter cells.

In our approach to detect CTCs using standard flow cytometry, we utilized MDA-MB-436 cells as a model for CTCs derived from TNBC tumors, labeling them with HAL^50,54,55^. HAL inside these cells converts into the fluorescent compound, PpIX, which emits strong fluorescence in mitochondria, especially in cancer cells due to their slower conversion of PpIX to heme^55^. Typically, the low concentration of PpIX in sparse CTCs results in insufficient signals for flow cytometry. However, we significantly boosted the signal by employing exosome-mediated homotypic targeting, further enhanced by modifying exosome surfaces with Tb^3+^ ions. HAL-loaded and Tb^3+^ ion-enhanced Exo-MDA exosomes effectively targeted CTC-like MDA-MB-436 cells, even in the presence of a high concentration of PBMCs. During this process, HAL was primarily contained within the Exo-MDA-Tb exosomes, preventing PpIX formation in the MDA-MB-436 cells. After centrifugation to remove unbound Exo-MDA-Tb exosomes, the cells were ultrasonicated to disrupt them completely. The contents released from these disrupted cells, including HAL, were then introduced into a pool of about 2×10^6^ MDA-MB-436 reporter cells (Fig. 5b). Here, HAL entered the cells through the PEPT1/2 channel and converted to PpIX, accumulating in their mitochondria (Fig. 5c-f). A 3-h incubation produced PpIX signals in roughly 1×10^4^ reporter cells for each CTC-like cell, significantly enhancing flow cytometric detection against a background noise level of approximately 1×10^4^ reporter cells. Notably, incubating 0.5 mM HAL with Exo-MDA exosomes for just one hour prior to their surface modification and capture by CTC-like cells resulted in efficient HAL loading into the exosomes and a substantial amplification of the final signal (Fig. 5g, h).

To evaluate the sensitivity, specificity, and LOD of the high-precision CTC detector, we conducted the ultrasensitive detection and enumeration of CTC-like cells in whole mouse blood using the cost-effective, single-color mode flow cytometer (Methods). Initially, we assessed sensitivity by counting CTC-like cells added to 1 mL of PBS (Extended Data Fig. 5a). When HAL was released from an Exo-MDA-Tb treated CTC-like cell into 2×10^6^ reporter cells, about 0.75% (or 1.5×10^4^ cells) exhibited an increased PpIX signal compared to the control without CTC-like cells (Extended Data Fig. 5b, c). A clear linear relationship emerged between the number of CTC-like cells and the rise in gated PpIX-positive reporter cells (Extended Data Fig. 5c). However, beyond 5 CTC-like cells, a nonlinear increase in reporter cell count occurred due to excessive HAL-loaded Exo-MDA in each cell, leading to saturation. We then introduced CTC-like cells into 1 mL of whole mouse blood to mimic samples with rare CTCs (Extended Data Fig. 5d). Flow cytometry showed outstanding detection capabilities, with signal intensity increases from 0.59% to 36.1% correlating with 1 to 20 spiked CTC-like cells in the blood (Fig. 6a, b and Extended Data Fig. 5e, f). Approximately 0.75% of reporter cells, which were equivalent to 1.5×10^4^ cells, showed a heightened PpIX signal compared to the control sample, which contained no CTC-like cells. This difference defined the baseline noise level for measuring the CTC signal (Fig. 6b). We verified a robust linear correlation when charting the number of CTC-like cells against the count of gated PpIX-labeled reporter cells, reflected by a linear fit with an R^2^ value of 0.9902 (Fig. 6b). Recognizing that mice with TNBC tumors might concurrently manifest secondary acute myeloid leukemia (AML) during their treatment course, we evaluated both the AML cell line, HL-60, and the non-TNBC BC cell line, MCF-7. The outcomes indicated elevated specificity, with negligible signal observed for exosomes from both HL-60 and MCF-7 cells (Fig. 6c). Fig. 6d emphasizes the unparalleled sensitivity and specificity of this method in recognizing CTC-like cells in a 1-mL whole blood sample, corroborated by ROC curves with impressive AUC values of 0.910, 0.965, and 0.990 for 1, 3, and 5 CTC-like cells. Furthermore, to determine the LOD, we examined 5-mL and 10-mL whole blood samples in addition to the 1-mL whole blood samples and consistently observed sensitive detection, discerning signals tied to a single CTC-like cell (Fig. 6e). The declining sensitivity trend is likely attributed to the increasing number of PBMCs, which may impede the efficient interaction between exosomes from CTC-like cells and reporter cells. Using the linear fit as our guide, we calculated the maximal volume of whole blood where the signal of a single CTC-like cell was equivalent to the background noise (i.e., control) as about 19 mL. These data suggest that our CTC detector is adept at amplifying signals, enabling the specific identification of just one CTC-like cell within over 10 mL of whole blood, amidst roughly 10^11^ erythrocytes and 10^8^ leukocytes.

**Fig. 6.**
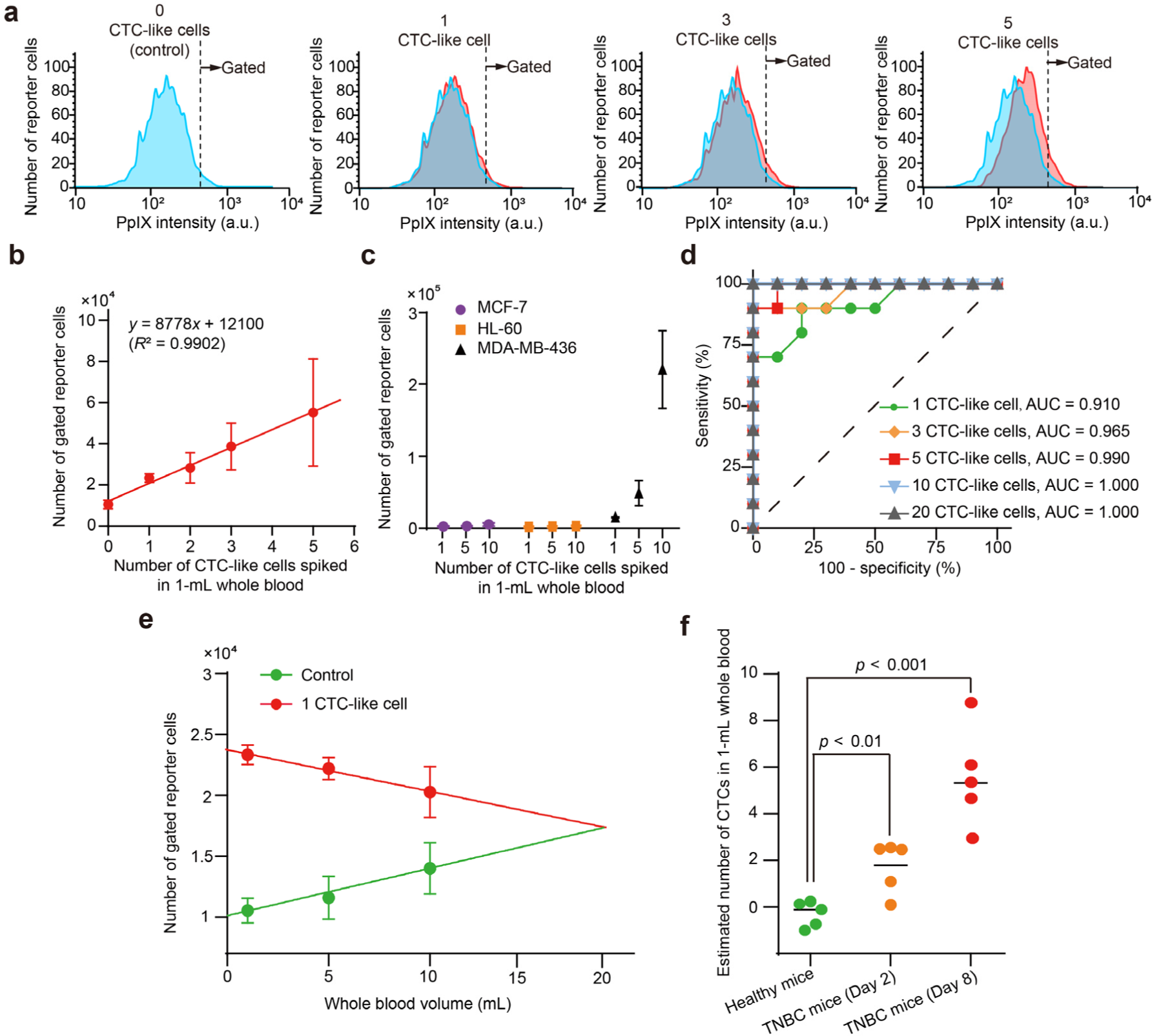
Results of extreme-precision CTC detection based on super homotypic targeting. **a**, Flow cytometry of reporter cells amplifying signals from varying numbers of CTC-like cells spiked in 1-mL mouse whole blood as shown in light red, in comparison with the control as shown in light blue (no CTC-like cells present). Each measurement used 2×10^6^ reporter cells. Dotted lines indicate gating thresholds for reporter cells. **b,** Number of gated reporter cells relative to different numbers of CTC-like cells spiked in 1-mL whole blood samples. Error bars represent standard deviations (n = 3). **c,** Number of gated reporter cells based on different CTC-like cells (MCF-7, HL-60, MDA-MB-436) spiked in 1-mL whole blood at varying numbers of spiked cells. **d,** ROC curves for detecting different numbers of CTC-like cells in 1-mL whole blood samples. **e,** Detection of a single CTC-like cell across varied whole blood sample volumes. The number of the gated reporter cells in the control (i.e., background noise) intersects with that from the whole blood sample containing a single CTC-like cell at around 18.9 mL. **f,** Number of detected TNBC-derived CTCs, estimated from the linear fit in **b**, from 1-mL whole blood samples taken on the 2nd and 8th days post-tumor implantation in mice. These results were compared with those from the control group (healthy mice) with n = 5 samples in each category. Each measurement used 2×10^6^ reporter cells. The p-values were obtained using a two-sided t-test to assess the statistical significance of the difference in the estimated number of CTCs between the “healthy mice” group and the “TNBC mice (Day 2)” group and between the “healthy mice” group and the “TNBC mice (Day 8)” group.

We used the fully evaluated CTC detector to directly detect CTCs originating from TNBC tumors in whole blood samples procured from TNBC-afflicted mice (Fig. 6f, Methods). Peripheral blood samples of 1 mL each were drawn on the 2nd and 8th days subsequent to the initiation of the TNBC xenograft in these mice. After the aforementioned preparation and CTC signal amplification steps, the samples were analyzed with the flow cytometer. The derived results revealed that a single run of flow cytometry, even with the affordable flow cytometer operating in single-color mode, differentiated between the whole blood samples of the control mice (n = 5) and the TNBC-afflicted mice (n = 5), based on the estimated count of CTCs derived from the linear fit in Fig. 6b with saturation-induced nonlinearity corrections (Extended Data Fig. 5f). By the 8th day, a range of 2-9 CTCs originating from TNBC tumors were detected within the blood of the TNBC-afflicted mice, whereas the blood samples from the control mice displayed an almost imperceptible estimated CTC count. This variance was statistically significant, evidenced by a low *p*-value (*p* < 0.001). Remarkably, as early as the 2nd day, a distinguishable disparity was observed between the two sets as emphasized by a robust *p*-value (*p* < 0.01), consistent with the ROC curve shown in Fig. 6d.

## Conclusions

In this study, we developed a method for super homotypic targeting by engineering exosome surfaces with lanthanide ions, demonstrating its efficacy in two precision detectors with exceptional performance. However, the scope of super homotypic targeting extends far beyond these applications, offering enormous potential across various biological and medical fields. In cancer therapy^5,10,13^, it could enable the precise targeting of tumor cells, minimizing damage to healthy tissue, which in turn improves treatment outcomes and lessens adverse effects. In tissue engineering^14,15^, it may foster more effective tissue regeneration and repair by promoting the growth and organization of specific cell types, crucial for healing wounds or replacing damaged organs. This approach could also significantly advance stem cell therapies by guiding stem cells to particular tissues or organs, thus improving their integration and effectiveness. A better grasp of homotypic interactions is expected to enhance our understanding of cellular communication, aiding in more accurate disease modeling and drug development. In immunotherapy^56^, it may enhance the targeting of immune cells, bolstering the body’s natural defenses against diseases. Additionally, this method has potential in combating drug resistance, a substantial challenge in treating chronic infections and cancer. Finally, super homotypic targeting aligns with the principles of personalized medicine, potentially enabling more precise tailoring of treatments to individual patient profiles.

## Methods

### Cell culture

MCF-7 and MDA-MB-436 cells were separately cultured in high glucose Dulbecco’s modified Eagle’s medium (DMEM) supplemented with 1% penicillin/streptomycin (PS) and 10% FBS. K562 cells were cultured in RPMI-1640 medium supplemented with 1% PS and 10% FBS. HL-60 cells were cultured in RPMI-1640 medium supplemented with 1% PS and 13% FBS. All cell lines were incubated at 37 °C in a 100% humidity atmosphere containing 5% CO_2_.

### Exosome isolation

Before isolating exosomes, MDA-MB-436 and MCF-7 cells were cultivated separately in DMEM supplemented with 1% PS and 10% exosome-free FBS for 48 h. Meanwhile, K562 cells were grown in RPMI-1640 medium enriched with 1% PS and 10% exosome-free FBS for the same duration. Once the MDA-MB-436 and MCF-7 cells reached about 70% confluence, their culture media were harvested. This was followed by a sequence of centrifugation steps on the 50 mL of collected media. Initially, cells and large debris were removed by centrifuging at 300 g for 10 min. The resulting supernatant was further centrifuged at 2,000 g for 10 min, and then at 10,000 g for 30 min. This was followed by an ultracentrifugation step at 100,000 g for 80 min. The sedimented exosomes, labeled as Exo-MDA for those from MDA-MB-436 cells and Exo-MCF7 for those from MCF-7 cells, were then resuspended in 500 μL 1× PBS and stored at -80 °C for subsequent experiments. For serum exosome extraction (Exo-BLD), either fetal bovine serum or serum from tumor-bearing mice (1.0 mL) was blended with 1.0 mL 1× PBS. Exo-BLD exosomes were then isolated using the identical centrifugation method applied for Exo-MDA and Exo-MCF7 exosomes. The Exo-BLD exosomes were finally resuspended in 1.0 mL 1× PBS and kept at -80 °C for later studies.

### Analysis of exosomes with western blotting (WB)

Exo-MDA, Exo-MCF7, and Exo-BLD exosomes were diluted to a concentration of approximately 4.0×10^7^ exosomes/mL). Proteins were extracted from the exosomes using RIPA buffer, supplemented with 1% protease and phosphatase inhibitor cocktail on ice. These proteins were then fractionated using SDS-PAGE and electrotransferred to nitrocellulose membranes. After blockading with 5% non-fat milk, the PVDF membranes were probed overnight at 4 °C with specific antibodies against TSG101, CD63, and Alix. This was followed by a 2-h incubation with the appropriate secondary antibodies at ambient temperature. Protein bands were finally visualized with ECL reagent.

### Chemicals and reagents

Hexaminolevulinate (HAL) hydrochloride (98%), Eu(NO_3_)_3_·6H_2_O (≥ 99%), 4′,6-diamidino-2-phenylindole dihydrochloride (DAPI) (≥ 95%), human serum albumin (HSA) (≥ 96%), R6G (≥ 95%), SA (≥ 98%), GSH (≥ 99%), proline (99%), cysteine (99%), L-histidine (≥99%), lysine (98%), glycine (≥ 99%), ATP (≥98%), glucose (98%), and PKH67 Green Fluorescent Cell Linker Kit were purchased from Aladdin Reagent (Shanghai) and Sigma-Aldrich (St. Louis, MO). Co(NO_3_)_2_·6H_2_O and TDA (≥ 98%) were purchased from Aladdin Reagent (Shanghai) and Fujifilm Wako Pure Chemical Corporation (Osaka), respectively. Other chemicals and reagents were obtained from commercial suppliers and used without further purification: RPMI-1640 medium and DMEM basic (1×) (Gibco), Penicillin-Streptomycin (Beyotime), exosome-depleted FBS (ThermoFisher).

### Preparation of (TDA)_x_(Co^2+^)_y_Eu^3+^

TDA (0.60 g) and Co(NO_3_)_2_·6H_2_O (0.89 g) were dissolved in 5.0 mL H_2_O, and gently shaken for 0.5 h. Then, 0.59 g Eu(NO_3_)_3_·6H_2_O dissolved in 5.0 mL H_2_O was added. The mixture was gently shaken for 1 h. The resulting (TDA)_x_(Co^2+^)_y_Eu^3+^ solution (0.15 g/mL) was stored at 4 °C.

### Preparation of TCT

TDA (0.69 g) and Co(NO_3_)_2_·6H_2_O (0.89 g) were dissolved in 5.0 mL H_2_O, and gently shaken for 1 h. Then, 0.43 g TbCl_3_·6H_2_O dissolved in 5.0 mL H_2_O was added. The mixture was gently shaken for 1 h. The resulting TCT solution (0.2 g/mL) was stored at 4°C.

### Detection of SA with (TDA)_x_(Co^2+)^_y_Eu^3+^

Various concentrations of SA solution, ranging from 0.032 mM to 8.08 mM (200 μL each), were combined with 200 μL of (TDA)_x_(Co^2+^)_y_Eu^3+^ solution (7.0 mg/mL) in water. Following a 5-min incubation, the fluorescence intensity of Eu^3+^ was recorded by a fluorescence spectrophotometer (RF-6000, Shimadzu). To evaluate the selectivity of SA detection, we examined potential interfering substances such as Na^+^, HPO_42−_, K^+^, glucose, GSH, glycine, HAS, and ATP. The conditions applied for SA detection were mirrored for this selectivity testing.

### Detection of SA with TCT

Various concentrations of SA solution, ranging from 0.5 mM to 10 mM (200 μL each), were combined with 200 μL of TCT solution (5.0 mg/mL) in water. Following a 5-min incubation, the fluorescence intensity of Tb^3+^ was recorded with a fluorescence spectrophotometer (RF-6000, Shimadzu). To evaluate the selectivity of SA detection, we examined potentially interfering substances such as lysine, proline, glycine, cysteine, L-histidine, GSH, and glucose. The conditions applied for SA detection were mirrored for this selectivity assessment.

### Engineering exosome surfaces with (TDA)_x_(Co^2+^)_y_Eu^3+^

For Exo-MDA-PKH, Exo-MCF7-PKH, and Exo-BLD-PKH exosome preparation: (i) The harvested exosomes from MDA-MB-436, MCF-7, and serum media were resuspended in 500 μL of diluent. (ii) Each exosome sample was stained with 0.8 μL PKH67 (MIDI67 -1KT) and agitated for 1 minute. (iii) The staining was finished by adding 10% exosome-depleted serum to remove excess PKH67. (iv) The mixtures were centrifuged at 100,000 g for 80 min. (v) Resultant Exo-MDA-PKH, Exo-MCF7-PKH, and Exo-BLD-PKH exosomes were then resuspended in 500 μL of 1× PBS. For Exo-MDA-Eu, Exo-MCF7-Eu, and Exo-BLD-Eu exosome preparation: (i) The extracted exosomes were suspended in 500 μL of 1× PBS. (ii) Each exosome preparation was stained with 0.8 μL of PKH67 (MIDI67 -1KT) or 0.5 μL of R6G (1.0 mg/mL) for 1 min. (iii) 8.0 μL of (TDA)_x_(Co^2+^)_y_Eu^3+^ solution was added to each mixture. (iv) After agitating the mixture for 1 min, it was centrifuged at 100,000 g for 80 min. (v) The resulting Exo-MDA-Eu, Exo-MCF7-Eu, or Exo-BLD-Eu exosomes were resuspended in 500 μL of 1× PBS.

### Engineering exosome surfaces with TCT

The surface of exosomes was modified with Tb^3+^ ions, which exhibit a sensitivity to SA comparable to that of Eu^3+^ (Extended Data Fig. 6a, b). This was done by adding TCT solution to exosome suspensions, obtained from cell culture media through ultracentrifugation. Exo-MDA exosomes from the TNBC cell line (MDA-MB-436) and Exo-MCF7 exosomes from the non-TNBC cell line (MCF-7) were altered in this way. After the TCT treatment, both Exo-MDA-Tb and Exo-MCF7-Tb exosomes showed increased size, due to aggregation prompted by the Tb^3+^ ions’ interaction with SA on exosomes. To maintain exosome sizes close to their natural range (90-150 nm), we used TCT at a concentration of about 0.1 mg/mL (Extended Data Fig. 6c). The rise in zeta potentials of these modified exosomes compared to their original forms indicates successful Tb^3+^ incorporation (Extended Data Fig. 6d). The greater changes in size and zeta potential in Exo-MDA-Tb exosomes, relative to Exo-MCF7-Tb exosomes, are likely due to the higher SA levels on Exo-MDA exosomes, reflecting the pronounced SA expression on MDA-MB-436 cell membranes (Fig. 2f) and resulting in more effective Tb^3+^ binding from TCT. Increased SA levels on cancer cells and their corresponding exosomes facilitate the attachment of Tb^3+^ ions to their membranes. Each Tb^3+^ ion interacts with two SA molecules, enhancing exosome-mediated homotypic targeting (Fig. 1a inset). For evaluating cellular exosome capture, Exo-MDA and Exo-MCF7 exosomes were pre-labeled with PKH67 prior to treatment with the TCT solution. We observed that MCF-7 cells took 5 h to capture both Exo-MCF7-Tb and Exo-MDA-Tb exosomes, while MDA-MB-436 cells showed a higher capture rate of Exo-MDA-Tb exosomes, as indicated by a significant increase in PKH67 fluorescence (Extended Data Fig. 7a-d). This greater capture by MDA-MB-436 cells is likely due to their higher SA content, improving exosome capture efficiency^20^. The use of Tb^3+^ ions was found to boost interactions between exosomes and cells, compared to unmodified exosomes (Extended Data Fig. 7e, f). Adding 0.8 µg/mL of TCT to the Exo-MDA-Tb exosome and MDA-MB-436 cell mixture resulted in even more pronounced PKH67 fluorescence, especially when the cells were in suspension (Extended Data Fig. 7d and 7g-i), demonstrating enhanced exosome delivery efficiency to target cells and reducing the incubation time from about 10 h to approximately 1 h. We further explored Exo-MDA-Tb’s targeted interaction with MDA-MB-436 cells. Exo-MDA and Exo-MDA-Tb exosomes, labeled with PKH67, were exposed to various cell lines including MDA-MB-436, B16, HepG2, and L02. After a 1-h TCT incubation, fluorescence microscopy showed MDA-MB-436 cells more effectively capturing Exo-MDA-Tb exosomes compared to the other cell lines (Extended Data Fig. 8). When MDA-MB-436 cells were introduced into mouse blood-derived PBMCs with varying concentrations of PKH67-labeled Exo-MDA-Tb exosomes (Extended Data Fig. 9a, b), LSCM revealed a specific affinity of Exo-MDA-Tb exosomes, at 1.6×10^6^ exosomes/mL, towards MDA-MB-436 cells, while mostly avoiding the PBMCs (Extended Data Fig. 9c).

### Protocol for engineering exosome surfaces with TCT

For surface-engineered Exo-MDA and Exo-MCF7 exosome preparation, we conducted the following steps: (i) The harvested exosomes from MDA-MB-436 and MCF-7 were resuspended in 500 μL 1× PBS. (ii) Each exosome sample was stained with 0.8 μL PKH67 (MIDI67 -1KT) and agitated for 1 min. (iii) 8.0 μL of TCT solution (0.1 g/mL) was added to each mixture. (iv) After agitating the mixture for 1 min, it was centrifuged at 100,000 g for 80 min. (v) The resulting Exo-MDA-Tb and Exo-MCF7-Tb exosomes were resuspended in 500 μL of 1× PBS. For HAL-labeled exosome preparation, we conducted the following steps: (i) The extracted exosomes from MDA-MB-436 cells were suspended in 500 μL of 1× PBS with 4.0×10^7^ exosomes/mL. (ii) Exo-MDA exosomes were stained with 100 μL of HAL (0.5 mM) for 1 h. (iii) 8.0 μL of TCT solution (0.1 g/mL) was added to the mixture. (iv) After agitating the mixture for 1 min, it was centrifuged at 100,000 g for 80 min. (v) Resultant Exo-MDA-Tb loaded HAL was resuspended in 500 μL of 1× PBS.

### LSCM of exosome capture by cancer cells in the exosome detection experiment

6×10^6^ MDA-MB-436 and 6×10^6^ MCF-7 cells were seeded in confocal dishes (3 cm in diameter) with 2 mL culture medium. Cells were treated with 20-40 μL of various exosome suspensions: Exo-MDA-PKH, Exo-MCF7-PKH, Exo-BLD-PKH, Exo-MDA-Eu, Exo-MCF7-Eu, or Exo-BLD-Eu exosomes. The suspensions of Exo-MDA-PKH, Exo-MCF7-PKH, Exo-MDA-Eu, and Exo-MCF7-Eu exosomes had concentrations of approximately 4.0×10^7^ exosomes/mL, while Exo-BLD-PKH and Exo-BLD-Eu exosomes were at about 3.8×10^8^ exosomes/mL. Cells were incubated with the exosomes for varying durations between 1 h and 40 h. After incubation, cells were rinsed with 1× PBS before undergoing confocal microscopy analysis.

### LSCM of exosome capture by cancer cells in the CTC detection experiment

MDA-MB-436 and MCF-7 cells were seeded in 6-well cell culture plates for 24 h. To validate selective exosome capture and improved homotypic targeting, cells were treated with 1.6×10^6^ exosomes/mL of different exosome suspensions: PKH67-labeled Exo-MDA-Tb or PKH67-labeled Exo-MCF7-Tb. Cells were incubated with the exosomes for varying durations between 1 h and 20 h. The cell nuclei were then stained with DAPI at 37 °C for 10 min. After incubation, the cells were rinsed with 1× PBS three times before undergoing LSCM analysis. Meanwhile, PBMCs, extracted from 1 mL of blood from healthy BALB/c nude mice, were spiked with 30 CTC-like cells (MDA-MB-436 cells). These cells, after a 3-h incubation in a culture medium with 1 mM HAL, were further treated with PKH67-labeled Exo-MDA-Tb in a culture medium for 1 h with 0.8 µg/mL TCT for varying concentrations of PKH67-labeled Exo-MDA-Tb ranging from 4.0×10^4^ exosomes/mL to 8.0×10^6^ exosomes/mL. The cells were washed three times with PBS and transferred onto a glass slide for verification of enhanced exosome capture using LSCM. The fluorescence signal of PpIX which was converted from HAL was utilized for validation of spiked CTC-like cells. This setup also confirmed the absence of Exo-MDA-Tb exosomes captured by PBMC components, including leukocytes. Furthermore, PBMCs extracted from 1 mL blood of healthy BALB/c nude mice were spiked with 1-30 CTC-like cells (MDA-MB-436 cells) and incubated with a culture medium containing 0.8 µg/mL TCT and 1.6×10^6^ exosomes/mL of HAL-loaded Exo-MDA-Tb for 1 h. Then, the uncaptured exosomes were discarded by centrifugation at 300 g. The mixture was resuspended in 0.5 mL of cell culture medium, disrupted by ultrasonication and introduced to around 2×10^6^ MDA-MB-436 cells as reporter cells seeded in a dish. The cells were then incubated for 3 h. Subsequently, the cells were washed three times with PBS and imaged using LSCM. Fluorescence intensity values shown in Fig. 3d, i, 5d, e and Extended Data Fig. 1h-j, 7d, f, h, and 9b were obtained by calculating the total fluorescence intensity from each cell and dividing it by the total number of cells (n = 30-50).

### In vivo imaging

Exo-MDA (500 μL in 1× PBS) was stained with DiR (2 μM) for 30 min. Subsequently, 8.0 μL of (TDA)_x_(Co^2+^)_y_Eu^3+^ solution was introduced to the mixture. Following a brief 1-min shake, it was centrifuged at 100,000 g for 80 min. The resulting Exo-MDA-Eu containing the DiR probe was resuspended in 500 μL of 1× PBS. Female BALB/c nude mice, 5 weeks old, were procured from Beijing Vital River Laboratory Animal Technology Co., Ltd. and were housed at the Pharmaceutical Animal Experimental Center of China Pharmaceutical University under specific pathogen-free (SPF) conditions, with care consistent with animal ethical standards. Upon developing the MDA-MB-436 tumor-bearing mouse model, the mice were categorized into three groups. They received either intratumoral or tail vein injections of 100 μL of DiR (100 nM), Exo-MDA (8.70 × 10^6^ exosomes/mL), or Exo-MDA-Eu (8.70 × 10^6^ exosomes/mL). Fluorescence imaging at intervals of 2, 4, 6, 8, and 24 h after injection was performed using the ABLX6 small animal imaging system (Tanon). At the experiment’s conclusion, 24 h after injection, the mice were humanely euthanized. Their organs including the heart, liver, spleen, lungs, kidneys, and tumors were immediately imaged with the same system.

### Preparation of TNBC-afflicted mice

BALB/c nude mice (female, obtained from Beijing Vital River Laboratory Animal Technology Co., Ltd.) were housed at the Pharmaceutical Animal Experimental Center at China Pharmaceutical University under specific pathogen-free (SPF) conditions and received care in accordance with ethical requirements for animal welfare (No. 220201299). The mice were injected with 1×10^6^ MDA-MB-436 cells into the mammary fat pad to induce the TNBC xenograft and facilitate the spontaneous generation of CTCs. All mice were randomized and their selection for injection or blood collection was conducted blindly.

### Extraction of PBMCs and preparation of CTC-like cells

To prepare PBMCs, 2 mL of lymphocyte separation medium was introduced into a tube, with 1 mL of mouse anticoagulant blood gently layered on top. Following a centrifugation at 500 g for 30 min, the mononuclear cell layer was carefully collected. This layer underwent another centrifugation to isolate the PBMCs, followed by two washes with DMEM culture medium. The resulting PBMCs, obtained from 1 mL of blood, were combined with suspended MDA-MB-436 cells in DMEM supplemented with 10% exosome-free FBS and 1% PS. 8.0 μL of TCT solution (0.1 g/mL) with 1.6×10^6^ exosomes/mL of Exo-MDA-Tb was added to the mixture and incubated for 1 h. The cells were washed with 1× PBS three times before imaging using LSCM.

### Flow cytometry

The BD FACSCelesta was used for all flow-cytometric measurements. The fluidic system of this instrument encompasses a sample injection module, sheath fluid reservoir, and a series of tubing and valves that facilitate smooth cell movement. It is equipped with multiple lasers (405 nm, 488 nm, 640 nm, 561 nm) and detectors, enabling simultaneous measurement of various parameters and ensuring the precise and accurate analysis of intricate cell populations. For analyzing host cells containing exosomes, the cell concentration was adjusted to 10^6^-10^7^ cells/mL by dilution with PBS. Once prepared, the sample was loaded into the instrument and analyzed using the appropriate laser and detector settings according to the fluorescent dye. The percentage of positive cells was determined by evaluating the fluorescence intensity of the cells.

### Imaging flow cytometry

Virtual-freezing fluorescence imaging (VIFFI) is an advanced optomechanical imaging method that addresses the common trade-offs between sensitivity, speed, and spatial resolution^39,40^. By “virtually freezing” the motion of moving cells on an image sensor, VIFFI achieves a 1,000-fold longer exposure time, allowing for microscopy-grade fluorescence image capture. Typically, VIFFI is integrated within flow-cytometric setups. Components of a VIFFI flow cytometer are as follows: (i) a flow-regulated microfluidic chip; (ii) a light-sheet optical excitation system, encompassing continuous-wave lasers (at 488 nm and 560 nm) for excitation, an excitation beam scanner, and a cylindrical lens; (iii) a scientific CMOS camera; and (iv) an optical imaging system featuring an objective lens, a 28-facet polygon scanner, situated in the Fourier plane of the fluorescence emitted by flowing cells, dual relay lens systems, each having four achromatic lenses, tailored to match the facet size while preventing aberrations, and a tube lens, which projects a wide-field fluorescence image of the flowing cells onto the camera. The working principle of VIFFI is as follows. The polygon scanner counteracts the motion of flowing cells on the camera. By rotating it against the flow direction at an angular speed matching the flow, it produces a sharp image of virtually stationary cells. However, solely countering motion is not enough for microscopy-grade imaging at high speeds, especially when considering factors like varying cell flow speeds and inevitable aberrations over a large field of view. To address this, a localized light-sheet excitation beam scans against the flow, covering the entire field of view but limiting local exposure to around 10 µs. This strategy reduces the demand for precise flow speed and minimizes system aberrations, leading to clearer images. The longer integration time on the camera is possible as long as the cell’s fluorescence remains within a single mirror facet of the polygon scanner. The scanner continues to rotate during the camera’s image data transfer. This ensures that its next facet aligns with the fluorescence’s Fourier plane as the camera’s subsequent frame begins. To prevent image gaps and handle randomly spaced cells, successive frames from the camera slightly overlap. This design ensures continuous capture at rates surpassing 1,000 frames per second, facilitating uninterrupted imaging of over 10 million cells at more than 10,000 events per second.

### Instrumentation

Fluorescence spectra of TCT and (TDA)_x_(Co^2+^)_y_Eu^3+^ for SA sensing were obtained using a Shimadzu RF-6000 instrument (Japan). Exosome isolation was facilitated by an L-70 Ultracentrifuge from Beckman Optima (USA). Morphological characterizations of the exosomes were conducted using a JEM-1400Flash TEM system from Japan. The Zetasizer Nano ZS90 instrument from Malvern Panalytical (England) was utilized to determine dynamic light scattering (DLS) and zeta potentials. Olympus’s laser-scanning confocal microscope (FV1200) was employed for cell imaging. ICP-MS readings were taken with an Agilent 7500 instrument (USA). Exosome concentrations were gauged using a ZetaView NTA PMX-120 device. Flow-cytometric analyses were executed using the FACSCelesta from BD Biosciences.

### Analysis of exosomes from cell lines with flow cytometry and imaging flow cytometry

We assessed Exo-MDA, Exo-MCF7, and Exo-BLD exosomes derived from cancer cell line culture mediums and serum using both the conventional flow cytometer (BD FACSCelesta) and the imaging flow cytometer (VIFFI flow cytometer). This was done to validate our method’s capability to distinguish Exo-MDA exosomes from other serum-or cancer-derived exosomes. Initially, we extracted the Exo-MDA, Exo-MCF7, and Exo-BLD exosomes from MDA-MB-436, MCF-7 culture mediums, and FBS, respectively. These exosomes were subsequently stained with PKH67 and modified by Eu^3+^, forming Exo-MDA-Eu, Exo-MCF7-Eu, and Exo-BLD-Eu exosomes. These stained exosomes, both individually and in mixtures, were then allowed to interact with host cells (MDA-MB-436 cells) over a 3-h incubation. Post incubation, exosome-marked host cells were detached using trypsin, thrice rinsed with 1× PBS, suspended in 1× PBS at a density of approximately 1.0×10^7^ cells/mL, and finally analyzed using both flow cytometric methods. For flow cytometry, host cells labeled with exosomes were enumerated based on PKH67 fluorescence. For imaging flow cytometry, the number of host cells labeled with exosomes was determined based on the identification of PKH67 fluorescence regions in the images of host cells.

### Analysis of exosomes from TNBC tumor cells with flow cytometry and imaging flow cytometry

To selectively analyze exosomes derived from TNBC, we executed the following steps: (i) Exosomes were extracted from 1.0 mL serum samples of healthy mice and TNBC-afflicted mice via centrifugation. (ii) These exosome mixtures were subsequently labeled with PKH67 and Eu^3+^. (iii) The stained exosomes were then incubated with host cells (MDA-MB-436 cells) for 3 h. (iv) Post incubation, cells were detached using trypsin, rinsed thrice with 1× PBS, and then suspended in 1× PBS at a density of approximately 1.0×10^7^ cells/mL. (v) These samples were subsequently analyzed using both the flow cytometer (BD FACSCelesta) and the imaging flow cytometer (VIFFI flow cytometer). The *p*-values for the results of both types of flow cytometry were calculated based on a two-sided *t*-test between healthy mice and TNBC-afflicted mice to assess the statistical significance of the difference between the two groups.

### Analysis of CTC-like cells with flow cytometry

We assessed CTC-like cells from the MDA-MB-436 cell line using the flow cytometer (BD FACSCelesta). The purpose of this assessment was to validate our method’s capability to detect single CTC-like cells from PBMCs. Initially, the known number of CTC-like cells (1-20 cells) were added to the PBMCs isolated from whole blood samples. The PBMCs with CTC-like cells were then incubated with a culture medium containing 0.8 µg/mL TCT and 1.6×10^6^ exosomes/mL of HAL-loaded Exo-MDA-Tb for 1 h. Following a centrifugation at 300 g for 3 min to discard uncaptured exosomes, the mixture was resuspended in 0.5 mL of cell culture medium. Subsequently, the cells were disrupted by ultrasonication and introduced to around 2×10^6^ MDA-MB-436 cells as reporter cells seeded in a dish. After 3-h incubation, the cells were detached using trypsin followed by trice washes with 1× PBS and suspension in 1× PBS at a density of approximately 1.0×10^7^ cells/mL. Finally, the reporter cells were enumerated using flow cytometry based on PpIX fluorescence.

### Analysis of CTCs derived from TNBC tumor cells with flow cytometry

To detect and enumerate CTCs derived from a TNBC tumor, we executed the following steps: (i) PBMCs were extracted from 1.0 mL whole blood samples of healthy mice and TNBC-afflicted mice via centrifugation. (ii) These cells were subsequently incubated with a culture medium containing 0.8 µg/mL TCT and 1.6×10^6^ exosomes/mL of HAL-loaded Exo-MDA-Tb for 1 h. (iii) The mixture was centrifuged at 300 g for 3 min to remove uncaptured exosomes. (iv) These samples were resuspended in 0.5 mL of cell culture medium, subjected to ultrasonication for cell disruption. (v) After ultrasonication, the contents were transferred to a dish containing approximately 2×10^6^ MDA-MB-436 cells as reporter cells. (vi) After a 3-h incubation, the cells were detached using trypsin and rinsed thrice with 1× PBS, and then suspended in 1× PBS at a density of approximately 1.0×10^7^ cells/mL. (vii) The reporter cell samples were subsequently analyzed using the flow cytometer (BD FACSCelesta). The *p*-values for the flow cytometry results were calculated based on a two-sided *t*-test between healthy mice and TNBC-afflicted mice to assess the statistical significance of the difference between the two groups.

## Data availability

The raw data obtained and the custom codes for all experiments are available on Mendeley Data (DOI: 10.17632/7w28np79vw.1). Any additional information required to reanalyze the data reported in this paper is available from the lead contact upon request.

## Acknowledgements

We thank the staff in the FACS Core Laboratory at the University of Tokyo. This work was mainly supported by JSPS Core-to-Core Program JPJSCCA20190007 (K.G.), JSPS KAKENHI JP19H05633 (K.G.), JP20H00317 (K.G.), JP21H05047 (S.S.), JP20KK0341 (S.S.), JP20H03252 (S.S.), JST START Program JPMJST2115 (M.S., K.G.), JST CREST Program JPMJCR1863 (S.S.), AMED 22ak0101163h0002 (M.S., K.G.), KAKETSUKEN (K.G.), White Rock Foundation (K.G.), G-7 Scholarship Foundation (M.S., K.G.), and China Scholarship Council 202107060015 (H.-S.W.).

## Author contributions

H.-S.W., T.D., Y.L., and K.G. conceived and refined the idea. H.-S.W. conducted the exosome surface modification and homotypic targeting enhancement experiments. X.-Y.H., Z.-W.Y, J.-L.S., Y.-Y.Y., F.-H.G., and J.-J.L. performed the animal experiments. H.-S.W., Y.L, Ya.Z., M.H., N.T.I., H.M., A.U., and H.G. conducted the flow cytometry and imaging flow cytometry experiments. H.-S.W., T.D., Yu.Z., Ya.Z. and K.G. analyzed and interpreted the experimental data. H.-S.W., T.D., Yu.Z., and K.G. wrote the paper. M.S., D.D.C., and S.S. provided valuable comments to improve the quality of the work. All authors participated in reviewing the paper. K.G. obtained most funding and supervised the work.

---

**Extended Data Fig. 1.**
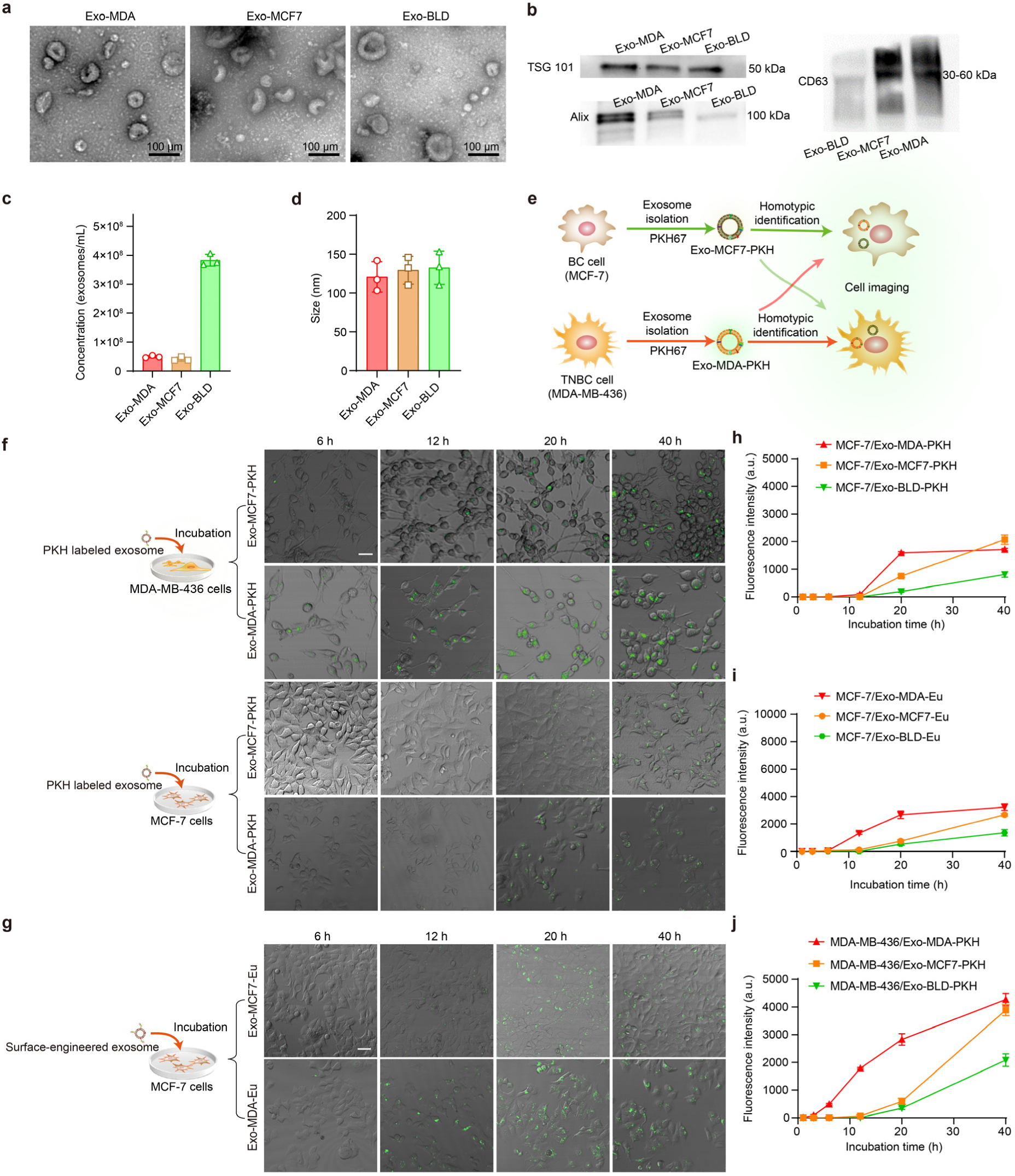
Cellular exosome-mediated homotypic targeting. **a**, Transmission electron microscope (TEM) images of exosomes. **b,** Western blotting (WB) analysis of exosome proteins: TSG101, Alix, and CD63. Exosomes were diluted to a concentration of approximately 4.0×10^7^ exosomes/mL. **c,** Concentrations of extracted exosomes. **d,** Hydrodynamic sizes of exosomes. **e,** Method for investigating the homotypic recognition of TNBC-and BC-derived exosomes. **f,** MDA-MB-436 cells incubated with Exo-MCF7-PKH or Exo-MDA-PKH. Scale bars: 30 μm. MCF-7 cells incubated with Exo-MCF7-PKH or Exo-MDA-PKH. Scale bars: 40 μm. The concentration of the Exo-MCF7-PKH or Exo-MDA-PKH suspension is ∼4.0×10^7^ exosomes/mL. The MDA-MB-436 and MCF-7 cells, seeded in confocal dishes, were incubated with 20 μL Exo-MCF7-PKH or Exo-MDA-PKH suspension for varying durations between 6 h and 40 h. Then, the cells were washed with 1× PBS and subsequently subjected to confocal microscopy analysis. **g,** MCF-7 cells incubated with Exo-MCF7-Eu or Exo-MDA-Eu for 6-40 h. The concentration of the Exo-MCF7-Eu or Exo-MDA-Eu suspension is ∼4.0×10^7^ exosomes/mL. The MCF-7 cells, seeded in confocal dishes, were incubated with the 20 μL Exo-MCF7-Eu or Exo-MDA-Eu suspension for varying durations between 6 h and 40 h. Then, the cells were washed with 1× PBS and subsequently subjected to confocal microscopy analysis. Scale bars: 40 μm. **h,** Relationship between the incubation time (1-40 h) and fluorescence intensity of MCF-7 cells incubated with Exo-MDA-PKH (20 μL), Exo-MCF7-PKH (20 μL), or Exo-BLD-PKH (20 μL). Error bars represent standard deviations (n = 3). **i,** Surface-engineered exosome capture for Exo-MCF7-Eu, Exo-MDA-Eu, and Exo-BLD-Eu by MCF-7 cells. Error bars represent standard deviations (n = 3). **j,** Relationship between the incubation time (1-40 h) and fluorescence intensity of MDA-MB-436 cells incubated with Exo-MDA-PKH (20 μL), Exo-MCF7-PKH (20 μL), or Exo-BLD-PKH (20 μL). Error bars represent standard deviations (n = 3).

**Extended Data Fig. 2.**
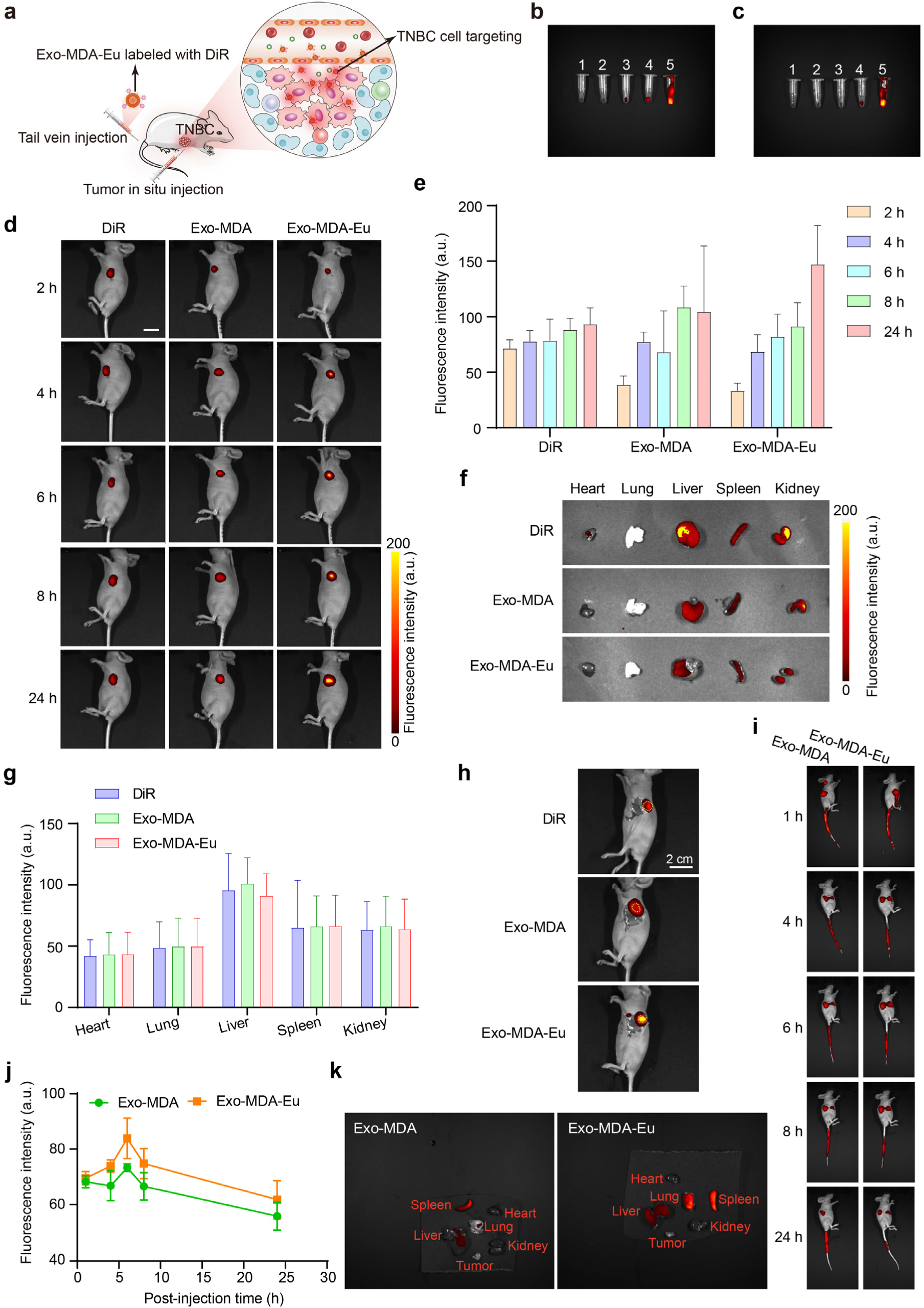
*In vivo* imaging of TNBC in mice. **a**, Schematic of *in vivo* TNBC tumor-targeting imaging using Exo-MDA-Eu (or Exo-MDA) through tail vein injection or tumor in situ injection. **b,** Exo-MDA and DiR samples with the following concentrations: 1: 8.70×10^5^ exosomes/mL of Exo-MDA; 2: 1.74×10^6^ exosomes/mL of Exo-MDA; 3: 4.35×10^6^ exosomes/mL of Exo-MDA; 4: 8.70×10^6^ exosomes/mL of Exo-MDA; 5: 100 nM of DiR. **c,** Exo-MDA-Eu and DiR samples with the following concentrations: 1: 8.70×10^5^ exosomes/mL of Exo-MDA-Eu; 2: 1.74×10^6^ exosomes/mL of Exo-MDA-Eu; 3: 4.35×10^6^ exosomes/mL of Exo-MDA-Eu; 4: 8.70×10^6^ exosomes/mL of Exo-MDA-Eu; 5: 100 nM of DiR. **d,** *In vivo* imaging of TNBC with DiR, Exo-MDA, and Exo-MDA-Eu introduced by intratumoral injections. Scale bar: 2 cm. **e,** Fluorescence intensity of *in vivo* imaging with DiR, Exo-MDA, and Exo-MDA-Eu. Error bars represent standard deviations (n = 3). **f, g,** Ex vivo fluorescence images (**f**) and fluorescence intensity values (**g**) of dissected organs from TNBC-afflicted mice after 24 h. Error bars represent standard deviations (n = 3). **h,** Fluorescence images of postoperative mice. The tumor tissue is highlighted by yellow circles. Exo-MDA-Eu exhibited the highest accumulation in the tumor region. **i,** *In vivo* TNBC images of TNBC-afflicted mice at 1-, 4-, 6-, 8-, and 24-h postinjection of Exo-MDA or Exo-MDA-Eu via tail vein. **j,** Fluorescence intensity values of TNBC-afflicted mice at 1-, 4-, 6-, 8-, and 24-h postinjection of Exo-MDA or Exo-MDA-Eu via tail vein. Error bars represent standard deviations (n = 3). **k,** Ex vivo fluorescence images of dissected organs from TNBC-afflicted mice.

**Extended Data Fig. 3.**
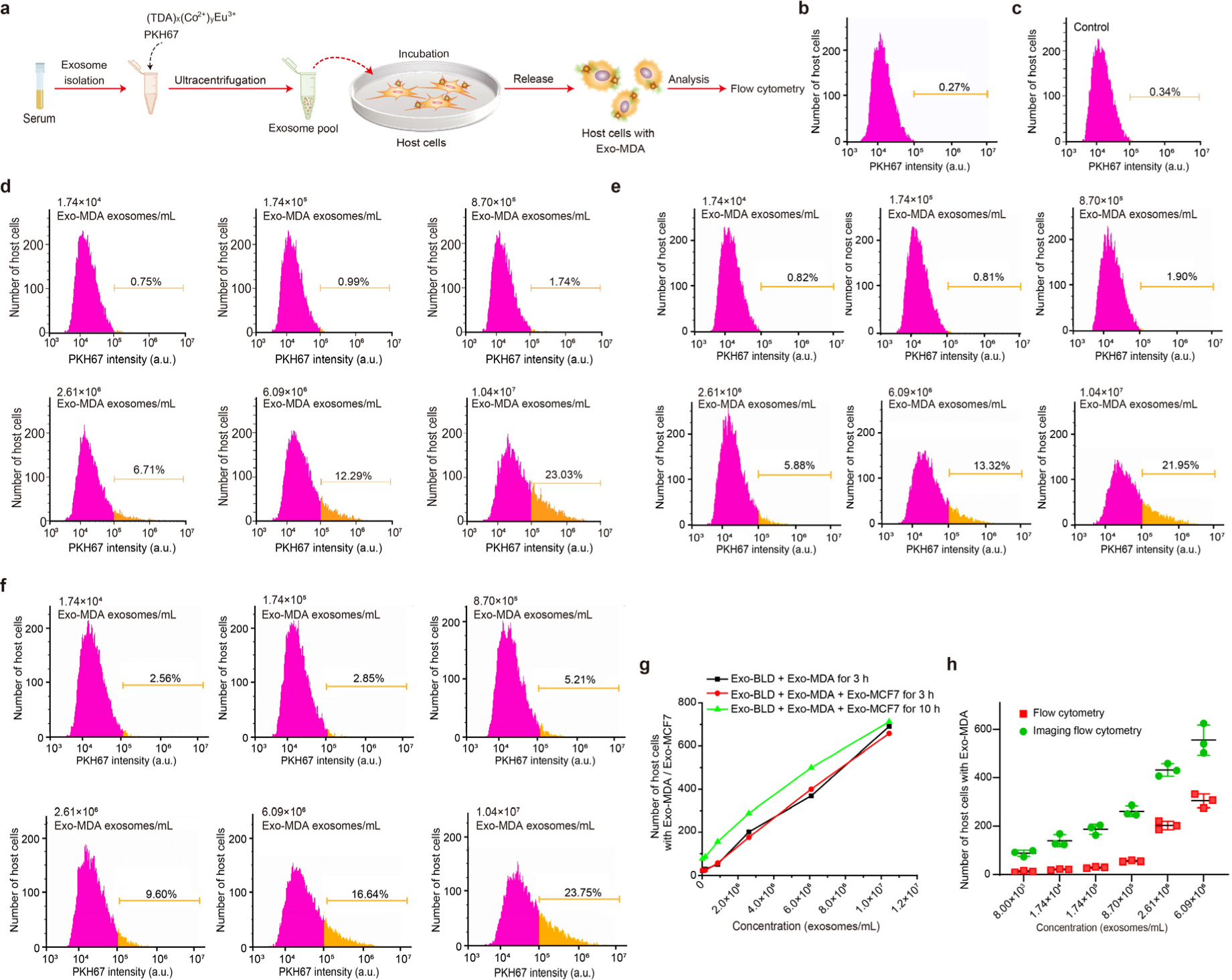
Flow-cytometric analysis for extreme-precision exosome detector based on super homotypic targeting. **a**, Schematic of the experiment. **b,** Number of host cells (pure MDA-MB-436 cells). MDA-MB-436 cells were suspended in 1× PBS and diluted to a concentration of 10^6^ – 10^7^ cells/mL. **c,** Number of host cells incubated in FBS (control). **d,** Flow cytometry of host cells incubated with different concentrations of Exo-MDA in serum (1 mL) containing about 4×10^8^ exosomes (Exo-BLD) derived from blood cells. **e,** Number of host cells incubated with Exo-MCF7 (2.0×10^5^ exosomes/mL) and different concentrations of Exo-MDA in serum and incubated with MDA-MB-436 cells for 3 h. **f,** Number of host cells incubated with Exo-MCF7 (2.0×10^5^ exosomes/mL) and different concentrations of Exo-MDA in serum and incubated with MDA-MB-436 cells for 10 h. **g,** Relationship between the flow-cytometric enumeration of host cells and exosome concentration for incubation with Exo-MDA for 3 h, incubation with a mixture of Exo-MDA and Exo-MCF7 for 3 h, and incubation with a mixture of Exo-MDA and Exo-MCF7 for 10 h. **h,** Number of host cells selectively capturing Exo-MDA exosomes at different spiked concentrations in serum, detected with flow cytometry and imaging flow cytometry. Error bars represent standard deviations (n = 3).

**Extended Data Fig. 4.**
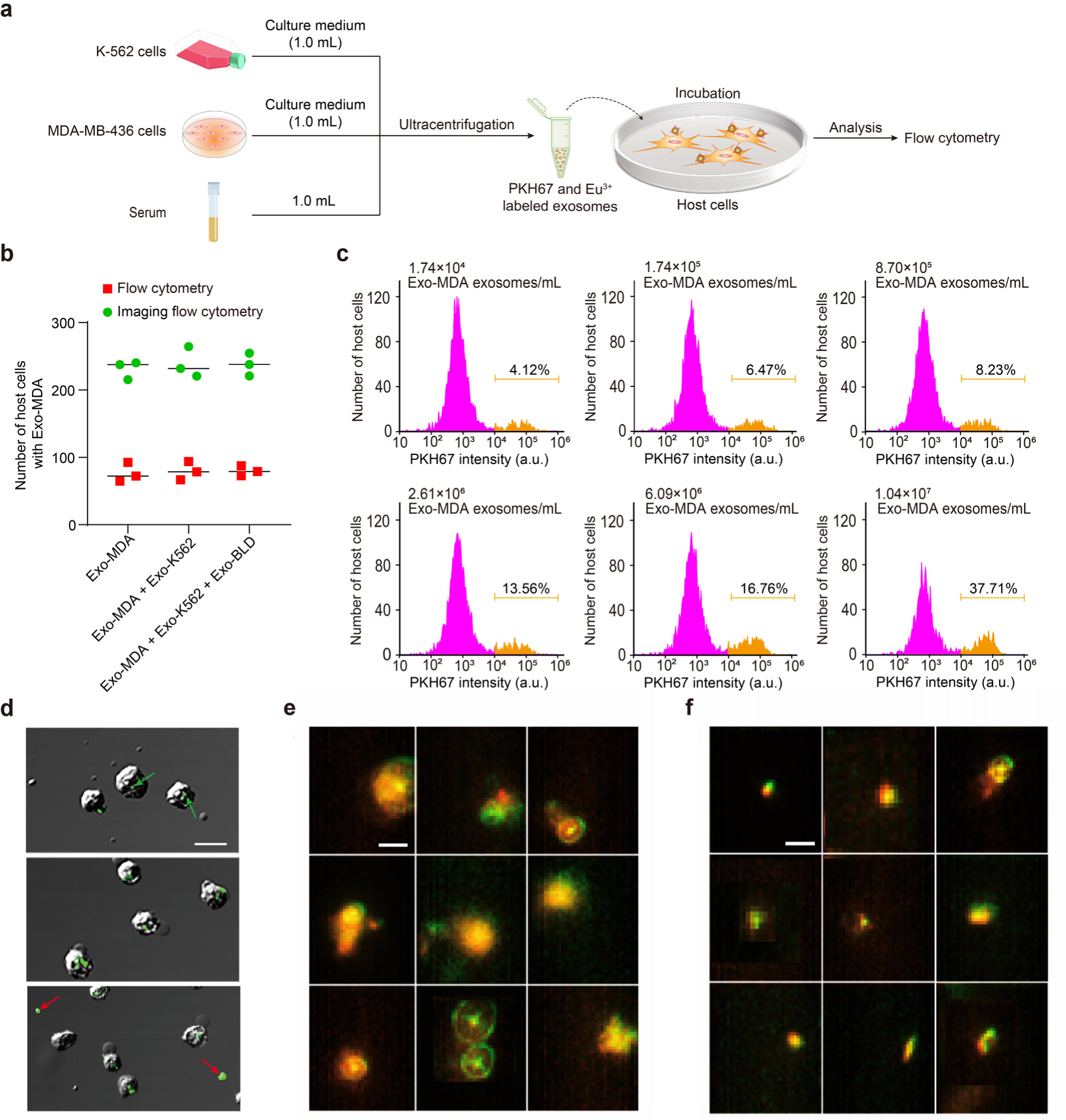
Selective detection of Exo-MDA exosomes in the presence of Exo-K562 exosomes and imaging flow cytometry of host cells. **a**, Schematic of the experiment for selective detection of exosomes derived from MDA-MB-436 cells (Exo-MDA) in the presence of exosomes derived from chronic myelogenous leukemia (Exo-K562). **b,** Number of host cells selectively capturing, isolating, and enriching Exo-MDA (1×10^7^ exosomes/mL) in the presence of Exo-K562 (1×10^7^ exosomes/mL) with and without Exo-BLD (4×10^8^ exosomes/mL) derived from serum samples of healthy mice. **c,** Number of host cells incubated with different concentrations of Exo-MDA, detected with imaging flow cytometry. **d,** Merged bright-field and LSCM images of MDA-MB-436 cells incubated with Exo-MDA prior to flow-cytometric analysis. Green arrows indicate cells labeled with a minimal capture number of exosomes. Red arrows indicate debris resulting in false positive events. Scale bar: 30 μm. **e, f,** Representative images of host cells labeled with Exo-MDA-Eu (green) and Exo-MCF7-Eu (red) with typical false positive (**e**) and positive (**f**) events, acquired with imaging flow cytometry. The Exo-MDA-Eu was labeled with PKH67 (green) while Exo-MCF7-Eu was labeled with R6G (red). The mixture of Exo-MDA-Eu (green) and Exo-MCF7-Eu (red) was incubated with the MDA-MB-436 cells for 3 h, and then analyzed with imaging flow cytometry. Scale bars: 20 μm.

**Extended Data Fig. 5.**
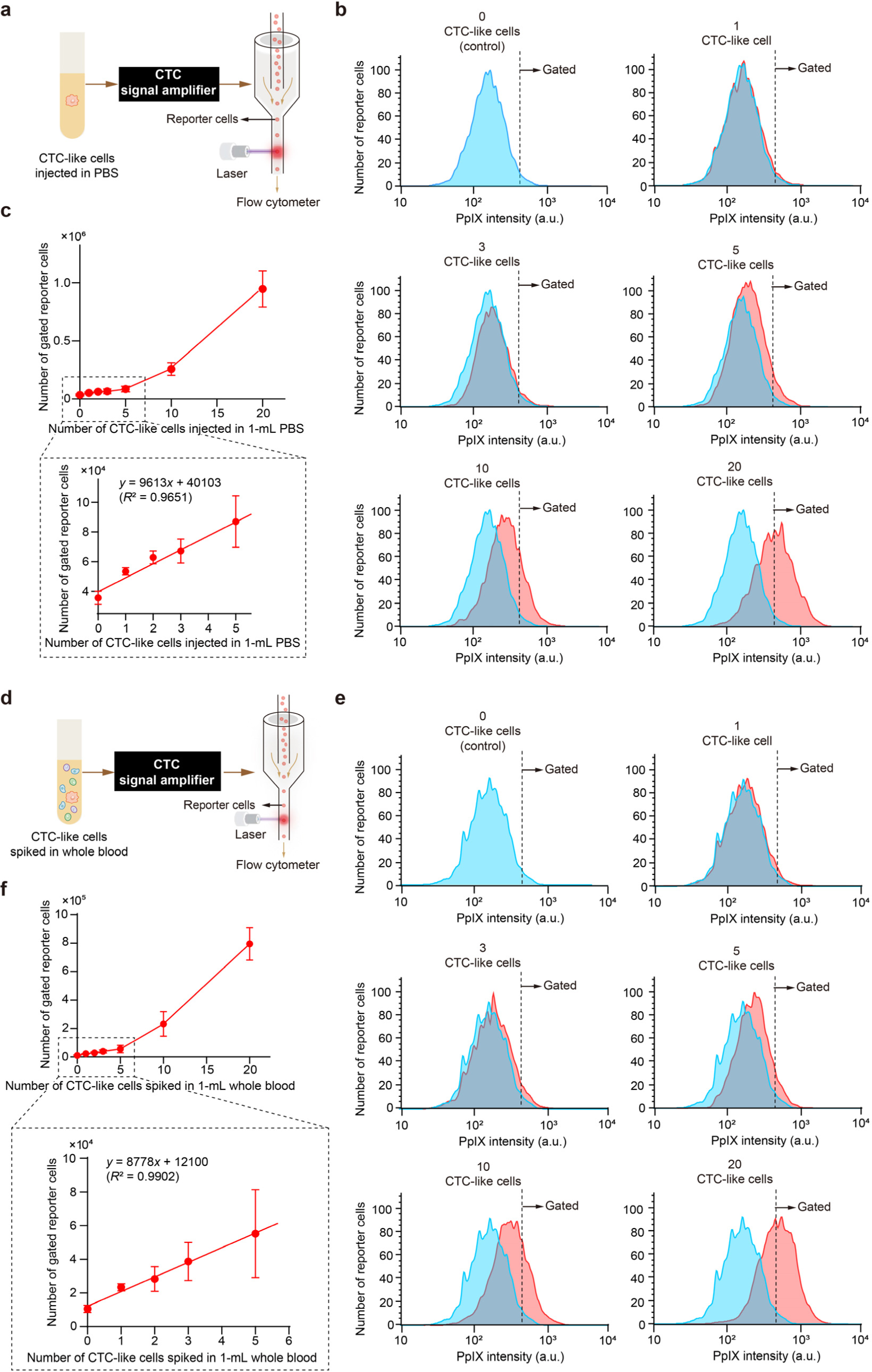
Spike-and-recovery validation of CTC-like cells with flow cytometry. **a**, Schematic of the experiment. CTC-like cells were injected into 1 mL PBS and prepared for flow-cytometric evaluation. **b,** Flow cytometry of reporter cells amplifying signals from varying numbers of CTC-like cells injected in 1-mL PBS, in comparison with the control (no CTC-like cells present). Each measurement used 2×10^6^ reporter cells. Dotted lines indicate gating thresholds for reporter cells. **c,** Number of gated reporter cells relative to different numbers of CTC-like cells injected in 1-mL PBS. Error bars represent standard deviations (n = 3). The increase in the error bar size with the increasing number of CTC-like cells is due to the increasing concentration of HAL in reporter cells, following a Poisson distribution. The reason why both the slope of the linear fit and the background noise are higher than those in Fig. 6b and Extended Data Fig. 5f is the absence of PBMCs that may capture HAL-loaded exosomes. **d,** Schematic of the experiment. CTC-like cells were spiked into whole mouse blood samples (1 mL each) and prepared for flow-cytometric evaluation. **e,** Flow cytometry of reporter cells amplifying signals from varying numbers of CTC-like cells spiked in 1-mL mouse whole blood as shown in light red, in comparison with the control as shown in light blue (no CTC-like cells present). Each measurement used 2×10^6^ reporter cells. Dotted lines indicate gating thresholds for reporter cells. **f,** Number of gated reporter cells relative to different numbers of CTC-like cells spiked in 1-mL whole blood samples. Error bars represent standard deviations (n = 3). The increase in the error bar size with the increasing number of CTC-like cells is due to the increasing concentration of HAL in reporter cells, following a Poisson distribution.

**Extended Data Fig. 6.**
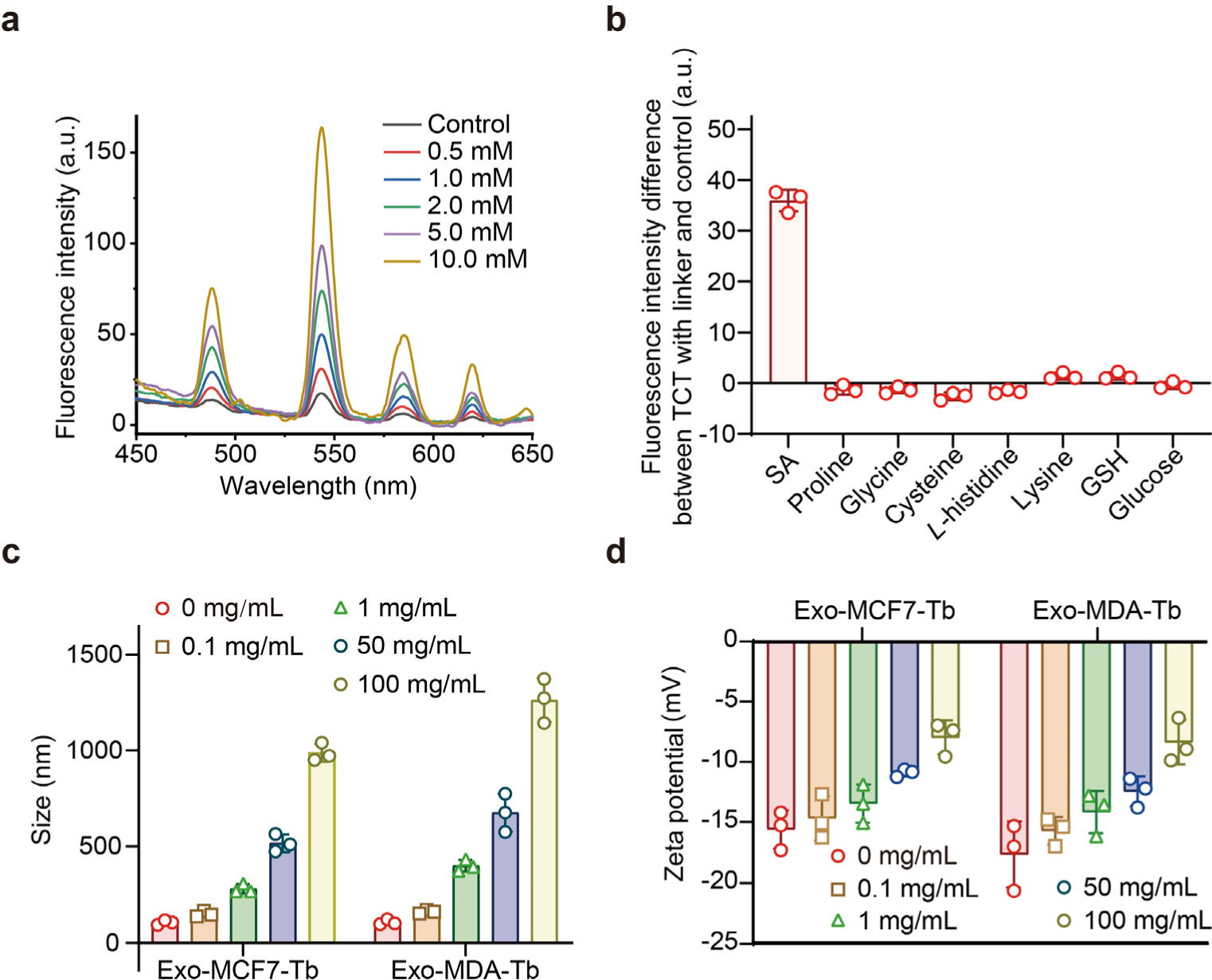
Exosome surface modification with Tb^3+^ ions for super homotypic targeting. **a**, Fluorescence spectra of TCT for sensing different concentrations of SA (excitation wavelength: 245 nm). **b,** Fluorescence intensity differences between TCT with different linkers and control. Error bars represent standard deviations (n = 3). **c,** Hydrodynamic sizes of surface-engineered exosomes with different TCT concentrations. Error bars represent standard deviations (n = 3). **d,** Zeta potentials of surface-engineered exosomes with different TCT concentrations. Error bars represent standard deviations (n = 3).

**Extended Data Fig. 7.**
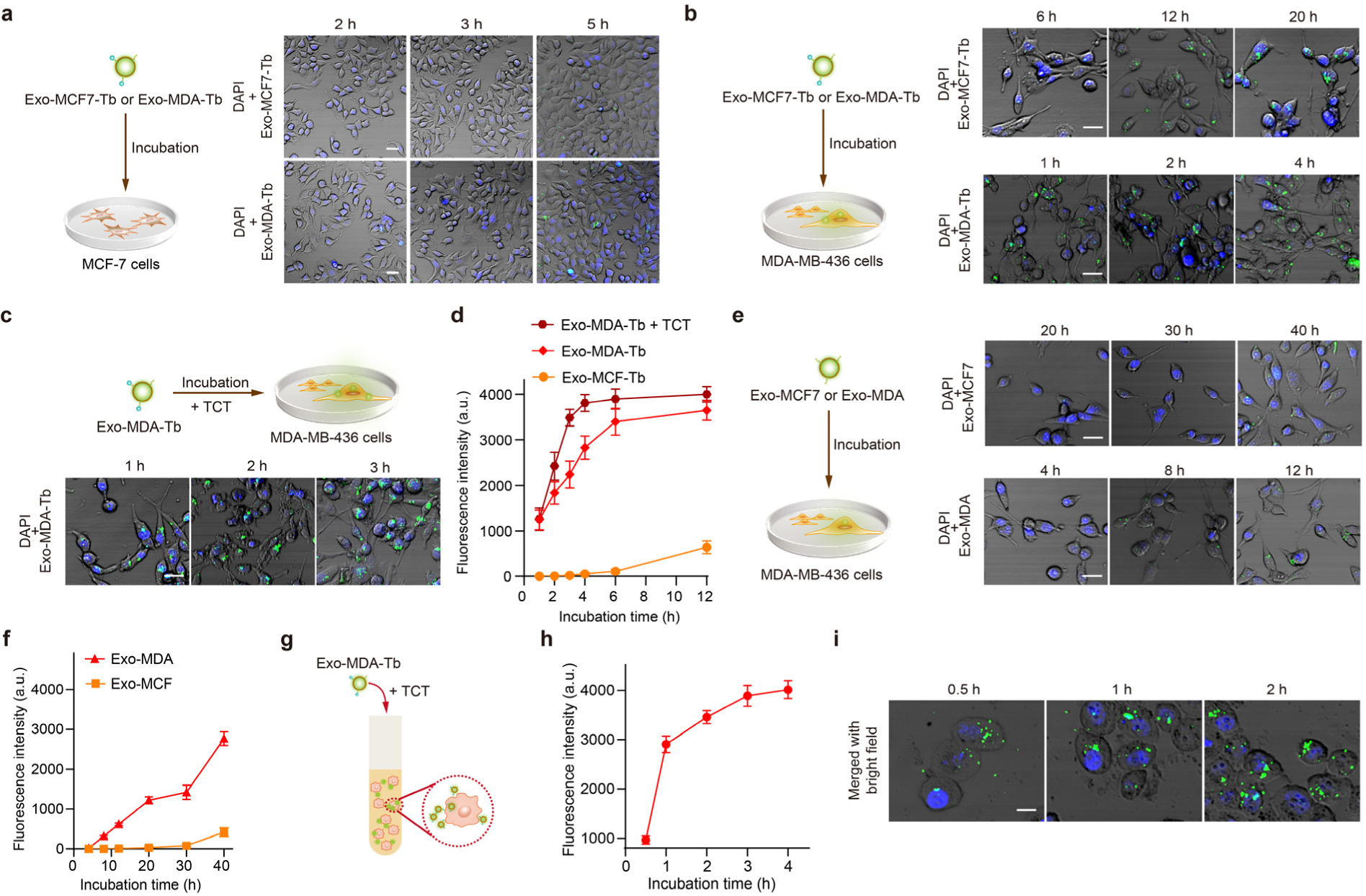
Selective capture and enrichment of exosomes modified with Tb^3+^ using MDA-MB-436 cells. **a**, Schematic of mixing MCF-7 cells with either Exo-MCF7-Tb or Exo-MDA-Tb, both labeled with PKH67, and LSCM images of MCF-7 cells after their mixture and incubation with Exo-MCF7-Tb and Exo-MDA-Tb for varying incubation durations (2-5 h). Scale bar: 30 μm. **b,** Schematic of mixing MDA-MB-436 cells with either Exo-MCF7-Tb or Exo-MDA-Tb, both labeled with PKH67, and LSCM images of MDA-MB-436 cells after their mixture and incubation with Exo-MCF7-Tb and Exo-MDA-Tb for varying incubation durations (6-20 h and 1-4 h). Scale bar: 20 μm. **c,** Schematic of mixing MDA-MB-436 cells with Exo-MDA-Tb labeled with PKH67 and TCT, and LSCM images of MDA-MB-436 cells after their mixture and incubation with Exo-MDA-Tb in the presence of TCT for varying incubation durations (1-3 h). Scale bar: 20 μm. **d,** Capture of surface-engineered exosomes by MDA-MB-436 cells for Exo-MDA-Tb in a culture medium with or without 0.8 µg/mL TCT and Exo-MCF7-Tb in a culture medium. Error bars represent standard deviations (n = 3). Fluorescence intensity values shown in this figure were obtained by calculating the total fluorescence intensity from each cell and dividing it by the total number of cells (n = 30-50). **e,** Schematic of mixing MDA-MB-436 cells with either Exo-MCF7 or Exo-MDA, both labeled with PKH67, and LSCM images of MDA-MB-436 cells after their mixture and incubation with Exo-MCF7 and Exo-MDA for varying incubation durations (20-40 h and 4-12 h). Scale bar: 20 μm. The concentration of exosomes was approximately 1.6×10^6^ exosomes/mL. **f,** Exosome capture by MDA-MB-436 cells for Exo-MDA and Exo-MCF7 in a culture medium. Error bars represent standard deviations (n = 3). The concentration of exosomes was approximately 1.6×10^6^ exosomes/mL. Fluorescence intensity values shown in this figure were obtained by calculating the total fluorescence intensity from each cell and dividing it by the total number of cells (n = 30-50). **g,** Schematic of mixing MDA-MB-436 cells with Exo-MDA-Tb labeled with PKH67 and TCT in suspension. **h,** Fluorescence intensity of suspended MDA-MB-436 cells after incubation with Exo-MDA-Tb in a culture medium with 0.8 µg/mL TCT for varying incubation durations (0.5-4 h). Error bars represent standard deviations (n = 3). Fluorescence intensity values shown in the figure were obtained by calculating the total fluorescence intensity from each cell and dividing it by the total number of cells (n = 30-50). **i,** LSCM images of suspended MDA-MB-436 cells incubated with Exo-MDA-Tb in a culture medium with 0.8 µg/mL TCT for varying incubation durations (0.5-2 h). Scale bar: 10 μm.

**Extended Data Fig. 8.**
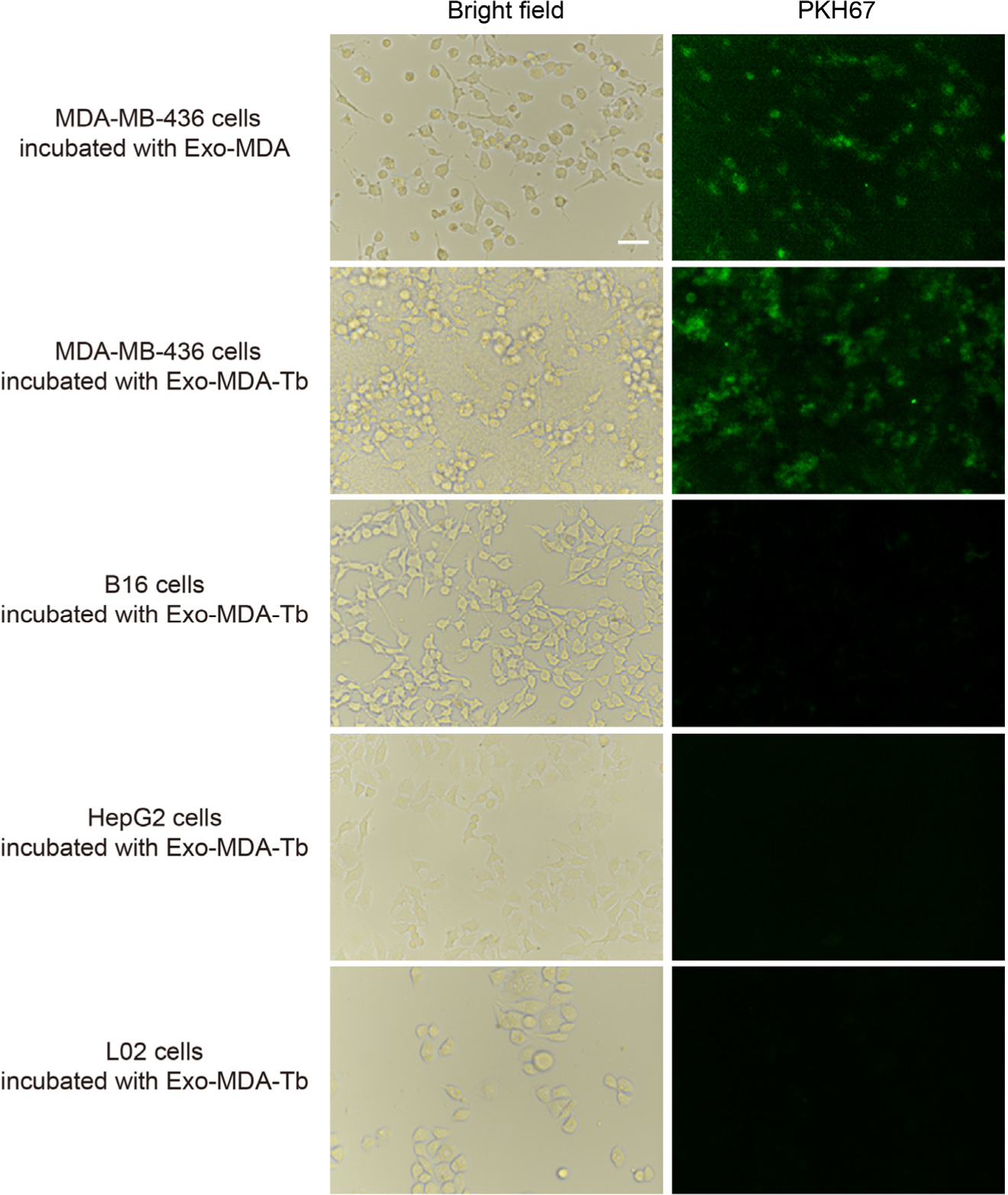
Verification of exosome-mediated homotypic targeting with different cells and exosomes. Inverted microscope images of different cell lines, including MDA-MB-436, B16, HepG2, and L02 cells, after their mixture and incubation with PKH67-labeled Exo-MDA or Exo-MDA-Tb (1.6×10^6^ exosomes/mL) in a culture medium with 0.8 µg/mL TCT for 1 h. Scale bar: 50 μm.

**Extended Data Fig. 9.**
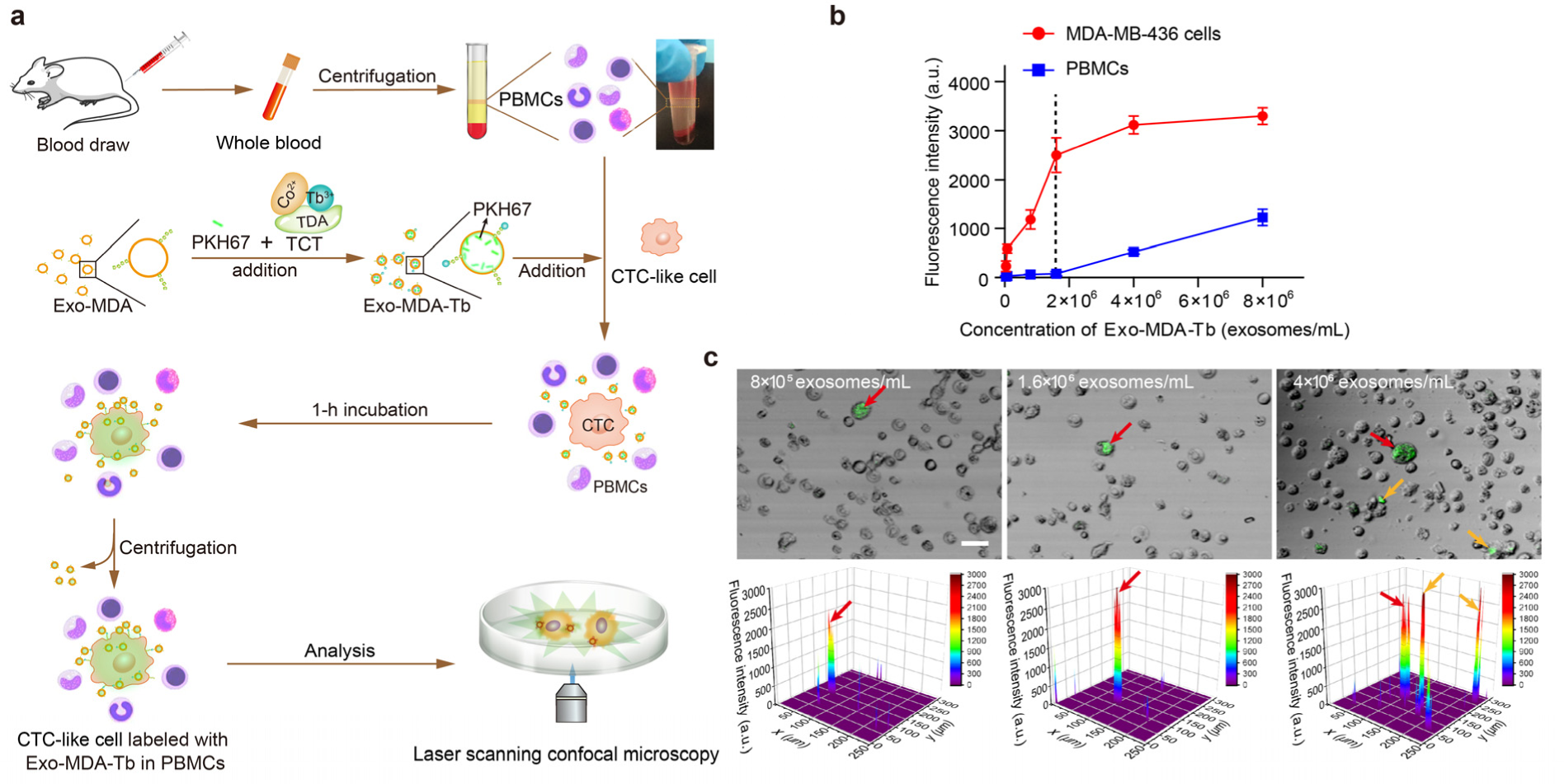
Verification of super homotypic targeting by surface engineering of exosomes with LSCM. **a**, Schematic of verifying super homotypic targeting by surface engineering of exosomes using LSCM. MDA-MB-436 cells were introduced into mouse blood-derived PBMCs as CTC-like cells before mixture and incubation with PKH67-labeled Exo-MDA-Tb. After centrifuging to discard uncaptured Exo-MDA-Tb, LSCM was performed to examine super homotypic targeting. **b,** Fluorescence intensities of suspended MDA-MB-436 cells and PBMCs incubated with varying concentrations of Exo-MDA-Tb in a culture medium with 0.8 µg/mL TCT for 1 h. The dotted line indicates the Exo-MDA-Tb concentration of 1.6×10^6^ exosomes/mL. Error bars represent standard deviations (n = 3). Fluorescence intensity values shown in the figure were obtained by calculating the total fluorescence intensity from each cell and dividing it by the total number of cells (n = 30-50). **c,** LSCM images and fluorescence intensities of suspended PBMCs with MDA-MB-436 cells after a 1-h incubation with Exo-MDA-Tb in a culture medium with 0.8 µg/mL TCT for varying concentrations of Exo-MDA-Tb. Red arrows indicate MDA-MB-436 cells with PKH67-labeled Exo-MDA-Tb and orange arrows depict PBMCs with PKH67-labled Exo-MDA-Tb. Scale bar: 30 μm

